# Community-Based Entomological Surveillance and Control of Vector-Borne Diseases: A Scoping Review

**DOI:** 10.1101/2024.09.13.612909

**Authors:** P. Eastman, T.S. Awolola, M. Yoshimizu, N. Govellla, P. Chaki, S. Zohdy

## Abstract

Community-based surveillance and control methods (CBMs) present opportunities to decentralize surveillance and control efforts while simultaneously enhancing community education, leadership, and participation in the fight against vector-borne diseases (VBDs). A scoping review was conducted to describe how CBMs are being utilized currently to combat malaria, dengue fever, Chagas disease, tick-borne diseases (TBDs) and other mosquito-borne diseases (MBD) exclusive of dengue and malaria, and to overall highlight key approaches, lessons learned, potential challenges, and recommendations. A total of 304 potential publications were identified among which 82 met the inclusion criteria. This scoping review highlighted the following benefits to CBMs: cost savings, increased sustainability, increased community knowledge, human behavior changes, increased surveillance coverage, ease in deployment, and the creation of larger, more diverse entomological datasets. Potential challenges highlighted include: participant retention and motivation, participant recruitment and incentives, continued governmental support, data quality, and collaboration with local municipal authorities. CBMs are commonly and successfully used in vector surveillance and control systems, but the chosen vector management method varies by vector-borne disease and region of the world. Additional research is needed to support the implementation of CBMs including cost-effectiveness studies and those studies with negative outcomes. Taken together, this scoping review highlights key aspects, potential challenges, and benefits of CBMs, and outlines potential future directions for incorporating CBMs into VBD control and elimination programming, and potential for community based integrated vector management (IVM) approaches.

## Introduction

According to the World Health Organization (WHO), vector-borne diseases (VBDs) account for more than 17% of all infectious diseases and cause more than 700,000 deaths annually, though many VBDs are preventable through control measures, particularly those that target the vectors that transmit them (WHO 2020). For example, malaria, dengue fever, Chagas disease, and more recently a suite of tick-borne diseases (TBDs) are of particular epidemiological importance due to their high burden, global populations at risk, and number of deaths annually. Malaria, a parasitic disease spread by *Anopheles* mosquitoes, caused an estimated 249 million cases in 2022 with around 608,000 deaths, the majority of which were in children under 5 years old in Africa (UNICEF 2024). Another mosquito-borne disease (MBD), dengue fever, is a viral disease spread by *Aedes* mosquitoes (*Aedes aegypti* and *Aedes albopictus*). Dengue fever causes an estimated 96 million cases per year in more than 129 countries which equates to almost half the world being at risk (WHO 2020, CDC 2023). Countries in Latin America are at greatest risk of Chagas disease, with around 6-7 million people expected to be infected annually with *Trypanosoma cruzi*, spread by triatomine bugs (WHO 2023). Lastly, the number of tick-borne infections is on the rise with the most common being Lyme disease, having around 476,000 cases per year in the United States and more than 200,000 per year in Europe (Marques 2021, CDC 2022).

There are many critical components for a successful fight against VBDs, such as case management, vector control, vector and disease surveillance, monitoring and evaluation, epidemiology, supply chain, and many more. Vector surveillance and control, two components that target the vector, are known to be some of the best preventative measures available against most VBDs. To interrupt transmission, vector surveillance is essential to ensure that public health vector control interventions are appropriately designed and effectively targeting vector populations. Additionally, vector control tools are developed based on understanding the biological and transmission characteristics of vectors including the dynamics of their behavior, all of which is gathered through surveillance (Killeen et al. 2017, Killeen et al. 2018).

Community-based methods (CBMs) are best distinguished from conventional or traditional vector surveillance and control methods due to their utilization of community members to assist or perform chosen interventions that would normally be conducted by a trained professional. In this scenario, ‘community members’ is defined as all individuals who live, work, and play in the chosen area, regardless of their leadership status, that are citizens born in the country. This term is inclusive of community-health workers but does exclude any residents already professionally trained in vector surveillance and control due to the focus being placed on training those without prior skill. CBMs can take on numerous variations depending on the disease system and vector at hand, but common variations examined in this review include community self-reporting of vectors, community active and passive surveillance, community working groups, and community health worker models. Conventional or traditional vector surveillance and control methods, though effective, are often most limited by a combination of cost, workforce constraints, and jurisdictional boundaries (Tokarz and Novak 2018, Little et al. 2019, Abrahan et al. 2021). Additionally, they tend to not directly involve the community in behavior change strategies for which they are the target population (Castro et al. 2012, Tana et al. 2012). As such, shifting from vertical to horizontal programming or towards CBMs, that include plans for effective community empowerment and mobilization are likely to provide a more sustainable approach to surveillance and control efforts (Castro et al. 2012, WHO 2017). The WHO Global Vector Control Response 2017-2030 recommends that vector surveillance and control interventions be tailored to the specific implementation area with the engagement of local authorities and communities to ensure that interventions are culturally appropriate and effective (WHO 2017). These recommendations have the potential to be framed through a community-led integrated vector management (IVM) approach- an evidence-based decision process to support a unified path for vector surveillance and control across disease systems. For effective and successful IVM and CBMs, social mobilization and capacity building should be the key elements prioritized (WHO 2012).

### Review Objectives

The objective of this scoping review is to utilize peer-reviewed literature to describe how community-based entomological surveillance and control is being utilized currently within the fields of malaria, dengue fever, Chagas disease, TBDs, and other MBD exclusive of dengue or malaria and to highlight 1) key approaches 2) lessons learned 3) potential challenges 4) recommendations and 5) present a case study of a comprehensive example from the literature. Additionally, based upon its popularity within specific disease systems and frequency in the literature, it became apparent to separately categorize research publications that were utilizing citizen science (CS); a common community-based approach. This CS section can be found in the discussion. The reason for the scoping review format is because of a current lack of research on the topic of CBMs especially in a condensed review. Both surveillance and control are covered in this review because surveillance is vital to ensuring efficient control methods are chosen and implemented successfully and control methods in conjunction with social mobilization are key elements to an IVM approach (Mutero et al. 2020). This paper reviews progress and lessons learned in community engagement in vector surveillance and control with a highlight on key areas where CBMs can improve. All the findings from this review can be found in a condensed framework format in supplemental document 1, which offers directions for focus on CBMs in each specific disease system.

## Methods

### Review Question

The review aimed to present how CBMs are being conducted globally across disease systems, contexts that drive and support community-based approaches, and lessons learned to create a final framework document consolidating all information for guidance (supp. Fig. 1).

**Figure 1.**
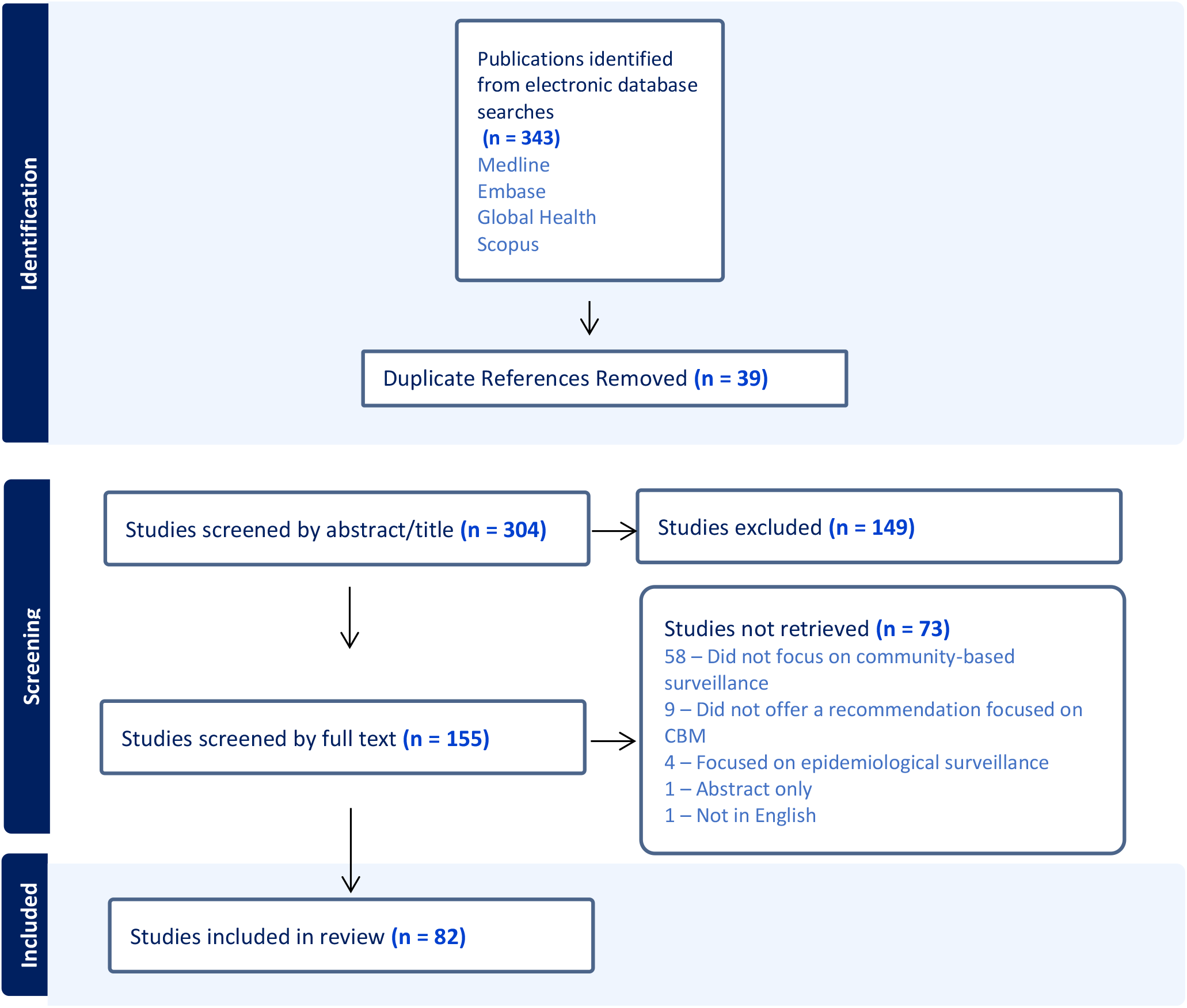
Scoping review PRISMA flowchart describing the scoping review process. PRISMA guidelines were used to structure the scoping review process. The scoping review process for this manuscript is outlined above describing the sources and literature reviewed for the process yielding the 82 studies included.

**Figure 2.**
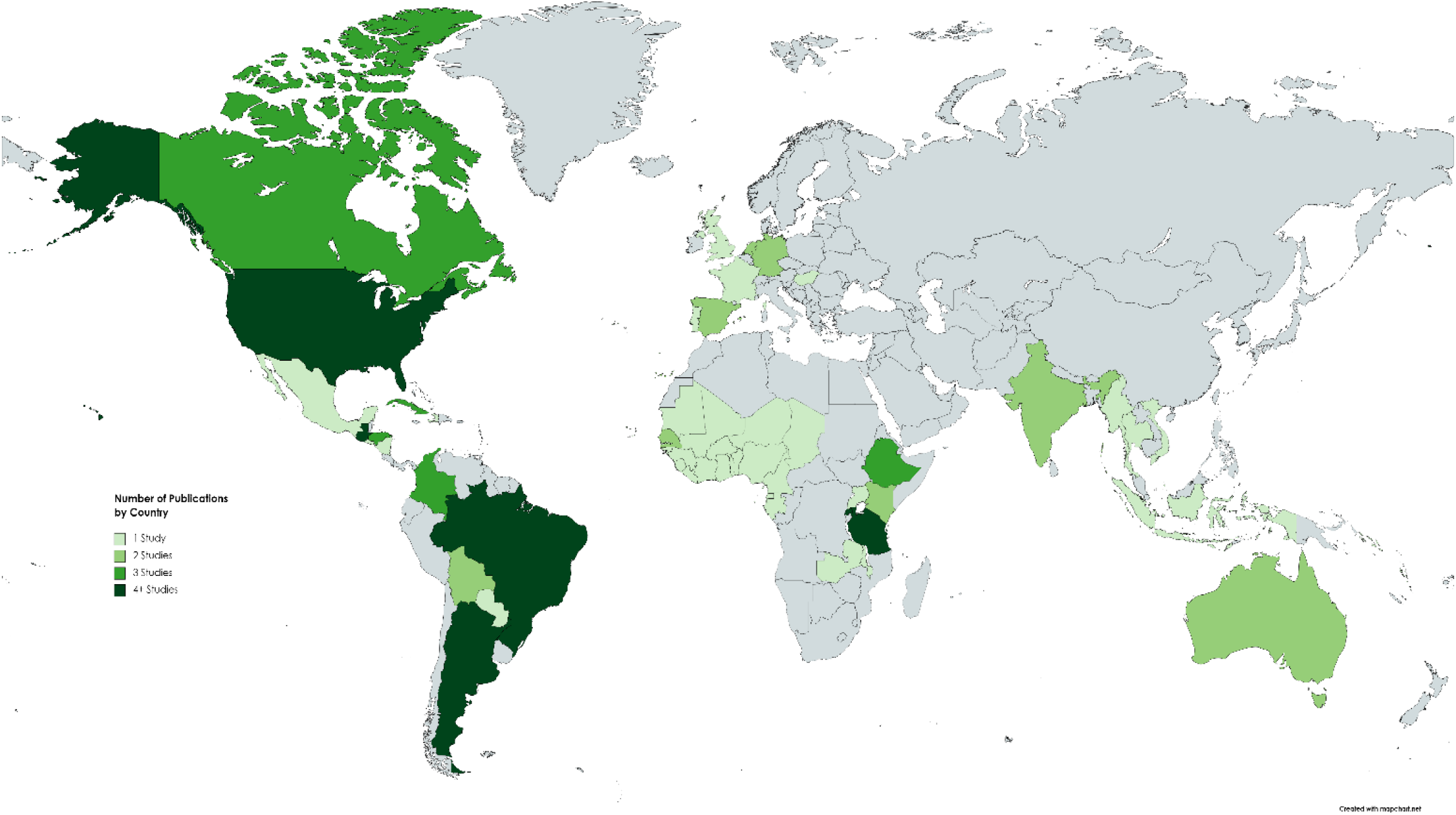
The global distribution of studies including in the scoping review. The distribution of community-based vector surveillance and control studies highlight that community-based activities occur globally with certain countries conducting the majority of studies in the region.

### Eligibility Criteria

Only publications in English were accepted, unless the article had an abstract given in both English and another language and an English full-text version of the article could be found. There were no limits on gender, age groups, or publication types. The first step of exclusions included: abstract only, publications focused on non-arthropod based vectors, and publications that focused only on epidemiological community surveillance and control or non-entomological surveillance and control. Finally, to gather all possible literature on the topic, articles that fit within at least one of the categories in Table 1 were included in the final selection.

**Table 1:**
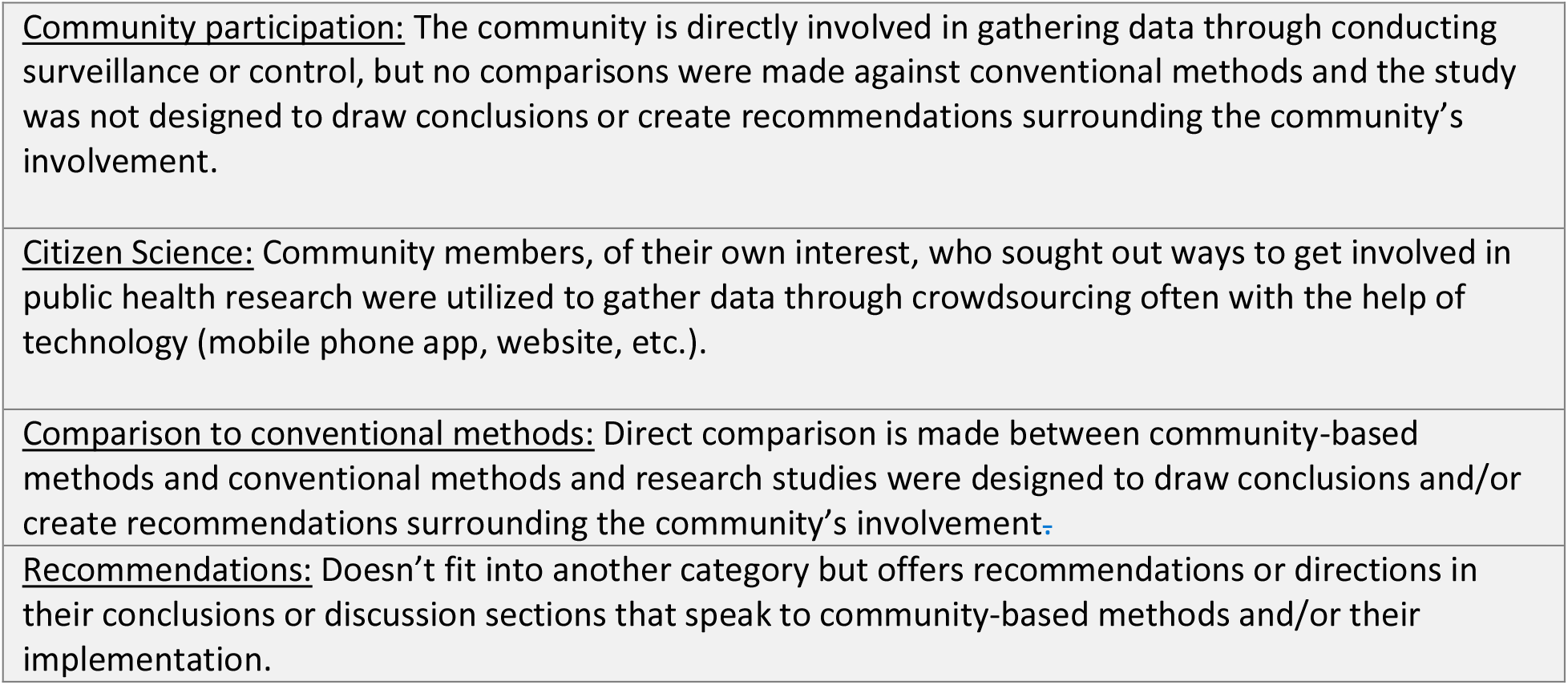
Publication Categories. To increase the number of articles included in the review and to identify any gaps, the categories in table 1 were utilized to make final inclusion decisions on all articles that passed the first round of inclusion/exclusion criteria.

|  |
| --- |
| <u>Community participation:</u> The community is directly involved in gathering data through conducting surveillance or control, but no comparisons were made against conventional methods and the study was not designed to draw conclusions or create recommendations surrounding the community's involvement. |
| <u>Citizen Science:</u> Community members, of their own interest, who sought out ways to get involved in public health research were utilized to gather data through crowdsourcing often with the help of technology (mobile phone app, website, etc.). |
| <u>Comparison to conventional methods:</u> Direct comparison is made between community-based methods and conventional methods and research studies were designed to draw conclusions and/or create recommendations surrounding the community's involvement. |
| <u>Recommendations:</u> Doesn't fit into another category but offers recommendations or directions in their conclusions or discussion sections that speak to community-based methods and/or their implementation. |

**Table 2:**
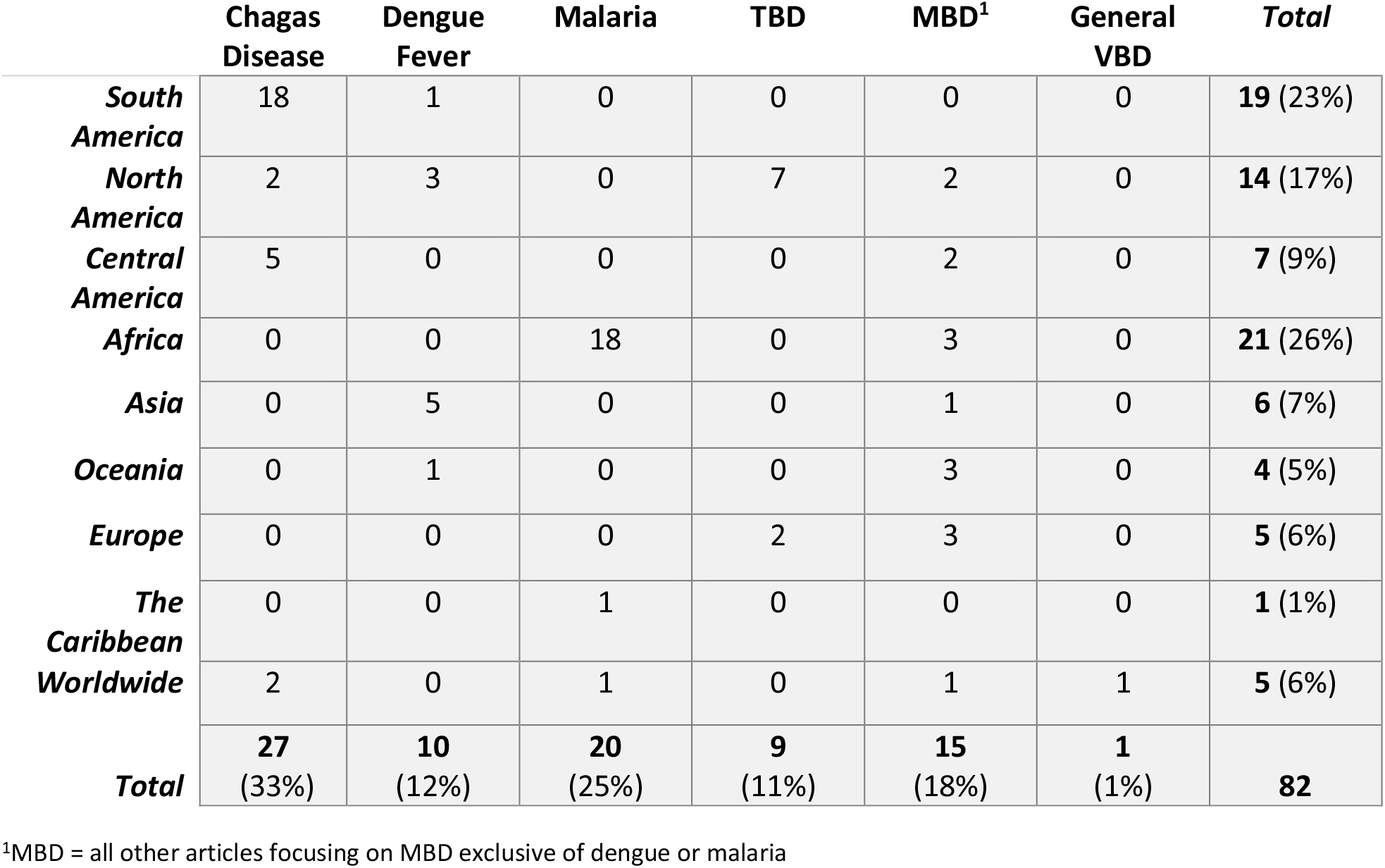
Final publications categorized by vector-borne disease and region of the world. Research was found across numerous regions and countries on the topic of CBMs. This table summarizes the breakdown of publications in each region by disease system.

|  | **Chagas**<br>**Disease** | **Dengue**<br>**Fever** | **Malaria** | **TBD** | **MBD<sup>1</sup>** | **General**<br>**VBD** | **Total** |
| --- | --- | --- | --- | --- | --- | --- | --- |
| **South**<br>**America** | 18 | 1 | 0 | 0 | 0 | 0 | **19 (23%)** |
| **North**<br>**America** | 2 | 3 | 0 | 7 | 2 | 0 | **14 (17%)** |
| **Central**<br>**America** | 5 | 0 | 0 | 0 | 2 | 0 | **7 (9%)** |
| **Africa** | 0 | 0 | 18 | 0 | 3 | 0 | **21 (26%)** |
| **Asia** | 0 | 5 | 0 | 0 | 1 | 0 | **6 (7%)** |
| **Oceania** | 0 | 1 | 0 | 0 | 3 | 0 | **4 (5%)** |
| **Europe** | 0 | 0 | 0 | 2 | 3 | 0 | **5 (6%)** |
| **The**<br>**Caribbean** | 0 | 0 | 1 | 0 | 0 | 0 | **1 (1%)** |
| **Worldwide** | 2 | 0 | 1 | 0 | 1 | 1 | **5 (6%)** |
| **Total** | **27**<br>**(33%)** | **10**<br>**(12%)** | **20**<br>**(25%)** | **9**<br>**(11%)** | **15**<br>**(18%)** | **1**<br>**(1%)** | **82** |

<sup>1</sup>MBD = all other articles focusing on MBD exclusive of dengue or malaria

### Search Strategy

Searches were limited to publications from 1980 to 2022, based on a Google ngrams (an online search engine that charts the frequencies of any set of search strings) searching the frequency of the phrase “Community-based surveillance” on the internet where the phrase peak started in 1980. Literature searches were conducted through databases Medline, Embase, Global Health, and Scopus. Key terms included in the search were: community-based entomological surveillance, vector-borne disease, vector control, IRS (Indoor Residual Spraying), ITNs (Insecticide-Treated Nets), LLINs (Long lasting insecticide treated nets), bed nets, community, surveillance, entomology, entomological monitoring, collection, decentralized, centralized, longitudinal, mosquito-borne, tick-borne, conventional, arbovirus, arboviral, arthropod, community-based control, citizen science, science, gold standard, and comparison.

Additionally, information on the search query can be found in supplemental figure 1. The review was undertaken from April to May 2022 using Endnote software to manage references and remove duplicates and Covidence software for management of the scoping review process. As an example, the search request used for Medline (Ovid) on 14 April 2022 was: ((community* ADJ5 surveillance) AND (entomological OR mosquito* OR tick* OR arthropod* OR vector-borne* OR vector control)).

### Selection of Studies

First, all retrieved publications were screened by title and abstract and were excluded if they were focused on non-arthropod based vectors and/or on epidemiological surveillance and control. The publications selected from title/abstract review underwent full-text review and were excluded from the final data extraction if they were abstract only and/or didn’t fit one of the publication categories in the above table. For both abstract and full-text screening, within the Covidence software (Melbourne, Australia) two independent reviewers selected the publications, and a third reviewer resolved differences. For data extraction, three independent reviewers conducted the extraction and compared results for a final consensus. Covidence software removed any duplicate articles during the initial importing step and any duplicates left over were found by reviewers, confirmed as duplicates, and removed.

### Study Characteristics and Data Extraction

An extraction template was created that allowed for the recording of the following information (supplemental figure 2): region, country, district, study site, disease, vector, methods used, aim of study, health metric used, focus of study, description of the focus of the study, limitations, and advantages to chosen focus, and overall conclusions and recommendations. Three contributors independently extracted data from the final 82 publications with results recorded in the extraction grid and any conflicts resolved amongst two contributors with the third contributor brought in when needed. The Preferred Reporting Items for Systematic Reviews and Meta-Analyses extension for scoping reviews (PRISMA-ScR) were used to present the review methods and the search results (Tricco et al. 2018). The scoping review was conducted following the framework proposed by the Joanna Briggs Institute guide for scoping reviews 2017 (JBI 2022). The protocol of the study was not registered but was developed a priori with fellow first authors.

## Results

The search strategy for the review identified 304 potentially relevant publications after duplicate references were removed. A final selection of 82 articles were identified that met the inclusion criteria for the review. (Preferred Reporting Items for Systematic Review and Meta-Analyses [PRISMA] flowchart, Figure 1).

The studies included in this review on global community-based vector surveillance activities are broken down by location of study and focal disease system in the study. The region with the largest number of publications is Africa (n = 21) and the majority of publications (n = 27) focus on Chagas disease. The countries with the highest number of publications included in this review by region are South America: Argentina & Brazil, North America: The United States of America, Central America: Guatemala, and Africa: Rwanda & Tanzania. To see the country breakdown in table form, please see Supplemental Table 1.

Of the four publication categories defined in the methods section for this review, the majority of publications included focused on community participation (37.3%), followed by citizen science (30.2%), comparison of CBMs to conventional methods (27.9%), and lastly, recommendations (4.6%). The results showed that publications within the ‘community participation’ category (56%) are primarily based in Central & South America with a focus on Chagas disease. Within the ‘comparison’ category, most publications (37%) are based in South America & Africa focusing on Chagas disease and malaria respectively. The region with the most publications utilizing ‘citizen science’ was North American (27%) with a broad focus on TBDs. Lastly, there were only 4 publications in the ‘recommendations’ category primarily focusing on malaria in Africa.

As stated in the methods, these publication categories were included as an additional categorization method to make final inclusion decisions on all articles and to increase the number of articles in the review per disease system due to the small amount of literature on the topic overall. These publication categories allowed the review to pinpoint areas where information is limited, and where gaps exist within each disease system to create areas for future research. For example, due to the majority of Chagas disease publications being within the ‘community participation’ category, we can see that while those research studies are actively using the community to conduct vector surveillance and control, the outcomes of that same research are not designed to address the benefits that involving the community had on the overall research. This is an important gap that future studies who plan on utilizing the community for data collection purposes should consider addressing.

To note, it is understood that the majority of CBMs in the review utilize vector surveillance management methods, but there are additional publications in the review that utilize vector control management methods especially within the dengue and other mosquito-borne disease systems (exclusive of malaria and dengue). These vector control management methods include larvicide application, environmental management, and many within the “other” category: mesocyclops application, home improvements, and fumigant canisters.

Table 3 shows the most common vector management methods utilized by each VBD, with vector management being defined as either a vector control or vector surveillance method. The results show that vector surveillance methods are very commonly used in Chagas disease studies while vector control methods are more commonly used in the study of MBDs such as dengue. Lastly, it can be seen that citizen science methods such as mobile phones were commonly used in the surveillance of TBDs and MBDs exclusive of malaria and dengue. Additionally, larval surveys, larvicide application, environmental management, and other are all vector management methods that are commonly used across disease systems. To see the breakdown of the “Other” category by disease see Supplemental Table 2.

**Table 3:**
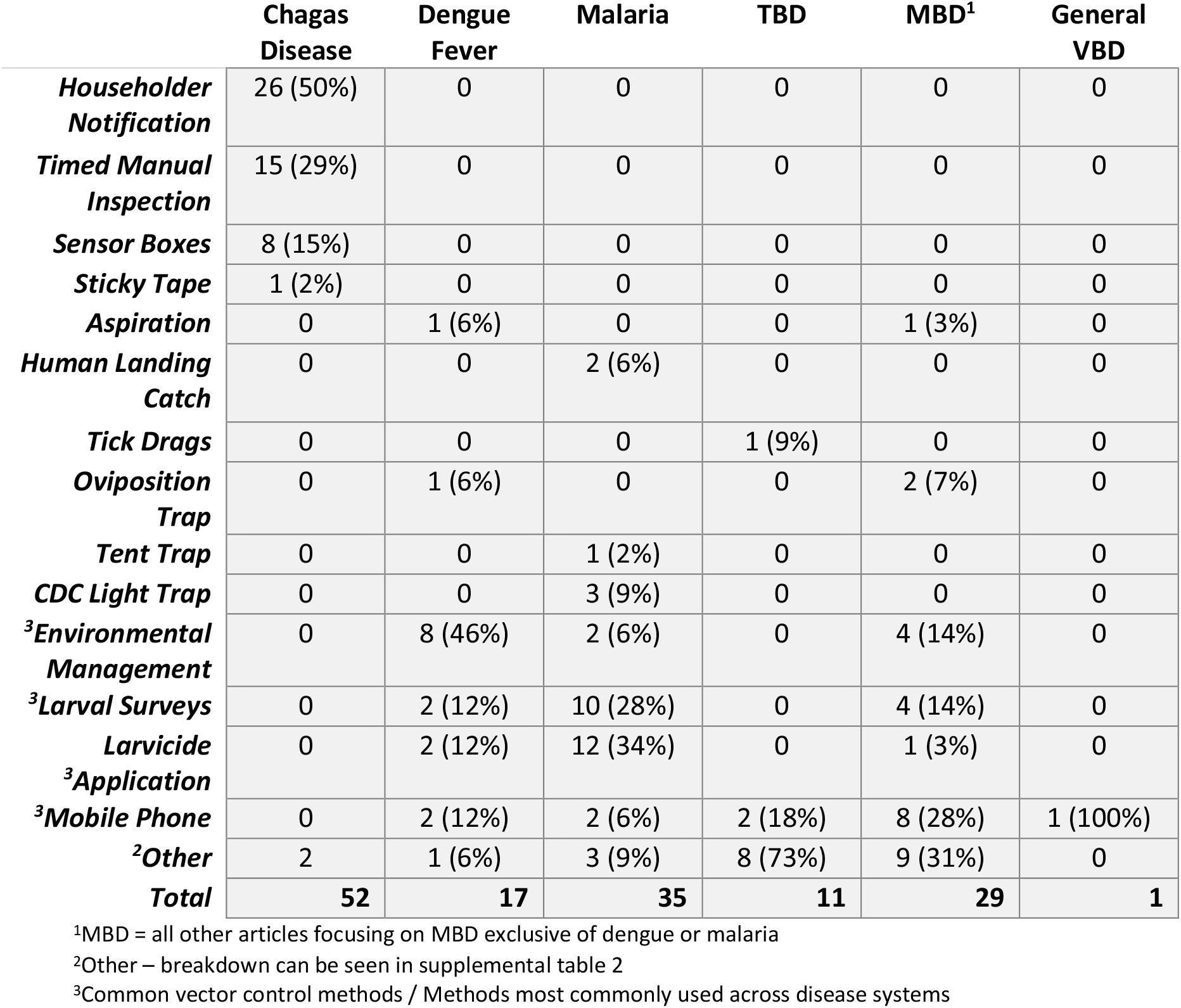
Final publications categorized by vector-borne disease and vector management method A variety of methods were used in the community-based vector surveillance and control. This table summarizes the collection and control methods used in the studies across disease and vector systems.

### Description of Findings of the scoping review

Every situation where VBDs are present is different and thus the vector control and surveillance solution implemented must be specific to the context of the disease, the vector, and the community. The sections below present the benefits and potential challenges for implementation of CBMs. For the development of guidance and ease of potential implementation, this discussion is broken down by disease system and split between those disease systems which do not frequently utilize citizen science (CS) and those that do. This decision to break down the discussion by disease system was made due to current vector surveillance and control systems primarily being developed and implemented within and for individual disease systems. Lastly, this discussion acknowledges the commonalities in the benefits and challenges of CBMs across all the disease systems presented and acknowledges the importance that the redundancy places on the most critical take-away findings.

#### Non-Citizen Science Focused Disease Systems

##### Chagas disease and householder notification and detection devices

In the context of Chagas disease, involving the community in entomological surveillance is noted as the best CBM and tool to help identify infestation and reinfestation of homes. This includes both active and passive surveillance methods. The main CBM recommended to reduce Chagas disease in the literature is householder notification (HN), which is an active surveillance method that involves householders actively collecting any suspected triatomine bugs in their home and sending them to local clinics or collection hubs for identification and/or analysis. HN has proven to be an effective method to increase collection days and specimens to create more robust datasets (Dumonteil et al. 2009, Ferro et al. 2018, Cavallo et al. 2018) and to increase empowerment and public knowledge (Yoshioka 2013, Parente et al. 2017, Rincon-Galvis et al. 2020). This method is noted as being easy to implement in areas with limited resources and is low-cost (Abad-France et al. 2011, Yoshioka et al. 2018). HN does have limitations which revolve mainly around householder motivation and interest in consistently collecting specimens and sending them to researchers or local health departments (Hashimoto and Yoshioka 2012, Abrahan et al. 2021, Leon et al. 2019), as well as poor capture technique and ability to identify larval stages (Rojas-Cortez et al. 2016). The main outcome found in the scoping review is that CBMs such as HN for Chagas disease are best introduced in areas with low infestation rates where frequent vertical control strategies are difficult. Methods such as HN offer an increase in collection coverage and thus can identify new foci or re-infestations quicker and more often than conventional methods which can also be beneficial in areas with frequent vertical control (Anonymous 1996, Abrahan et al. 2021). Due to the increase in surveillance coverage by HN, household insecticide spraying can be reduced to only the homes that need it, based on infestation reporting, thus reducing costs of current vertical programs (Rojas-Cortez et al. 2016, Abrahan et al. 2021). However, for HN to be most effective, both local health authorities and householders must be committed to providing a rapid response in reporting identified triatomines and responding to those households (Provecho et al. 2014, Hashimoto et al. 2015a, Cavallo et al. 2018, Abrahan et al. 2021). Other CBMs that householders can conduct include passive surveillance methods through detection devices such as sticky tape or sensor boxes. These devices relieve participants from having to actively looking for triatomines since they are designed to trap the specimens and any remnants of specimens such as excretion or urine. Sticky tape is cheap, can be easily administered by householders, and can be used to help make decisions as to whether certain communities should be targeted again for full-coverage insecticide spraying (Enriquez et al. 2020). But sticky tape was found to be unsuited for peri-domestic environments and is recommended only to be used in domestic ones (Enriquez et al. 2020, Abrahan et al. 2021). Sensor boxes are found to be cheaper and quicker at locating reinfestations though there can often be problems with installation and evaluation (Garcia-Zapata and Marsden 1993, Weeks et al. 2014). It is recommended that a combination method be utilized, one that involves both passive and active surveillance, such as sensor boxes or sticky tape and HN, since no one method offers 100% sensitivity in detecting triatomines (Abad-France et al. 2011, Weeks et al. 2014). It is shown that developing a targeted vertical attack phase, such as an insecticide spraying campaign, with the results from a horizontal/participatory phase, is the most beneficial in interrupting transmission (Cardinal et al. 2007, Hashimoto et al. 2015b, Yashioka et al. 2018, Cecere et al. 2019, Abrahan et al. 2021).

##### Dengue fever transmission and community-based environmental management

The literature has shown that unlike Chagas disease, which best utilize the community through entomological surveillance methods, dengue fever was found to utilize the community most often through vector control methods. The main CBM recommended for dengue fever targets the larval stage of mosquitoes through environmental management techniques such as source reduction conducted by community members (Nam et al. 2004, Toaliu and Taleo 2004, Sanchez et al. 2009, Vanlerberghe et al. 2010, Tana et al. 2012, Castro et al. 2012, Kittayapong et al. 2012). Source reduction and community empowerment can be achieved through the creation of working groups composed of community members who provide education on source reduction to households through door-to-door visits or community workshops (Vanlerberghe et al. 2010, Castro et al. 2012, Kittayapong et al. 2012, Wai et al. 2012, Parra et al. 2020). The creation of working groups increases the number of community members involved in the decision-making process of vector control programs, it helps create a shared sense of ownership on the local level which can increase program sustainability, and increase the level of knowledge regarding dengue and its vectors amongst the whole community (Nam et al. 2004, Vanlerberghe et al. 2010, Tana et al. 2012). Additional educational groups have been made utilizing student volunteers who bring home the lessons they have learned to protect their own communities (Parra et al. 2020). An additional benefit of the focus on environmental management is the intersectoral collaboration that happens between the health and sanitation sectors with source reduction interventions (Toaliu and Taleo 2004, Tana et al. 2012). Drawbacks of this method are that the creation of these working groups requires strong leadership, can often require greater investment at the beginning, that participants require effective interpersonal communication skills, and that they can be slower than vertical approaches in achieving goals (Tana et al. 2012, Kittayapong et al. 2012, Parra et al. 2020). Sustainability of these programs also relies on continued support from the government, community interest and ownership, and active participation (Sanchez et al. 2009, Kittayapong et al. 2012, Jongejan et al. 2019). Community empowerment and source reduction education can also be spread through educational materials such as posters, leaflets, pamphlets, radio or tv messages, trainings, speeches, and more. All educational materials are best distributed with the help of community leaders, health centers, city offices, and through the community working groups. These educational materials must be locally relevant and culturally acceptable to be effective (Sanchez et al. 2009, Vanlerberghe et al. 2010, Tana et al. 2012, Wai et al. 2012). It is recommended that specific biological or chemical larval control tools offered through vertical means, such as larvicide application, be blended with a horizontal program such as household managed source reduction and community empowerment through working groups to maintain effective coverage (Nam et al. 2004, Toaliu and Taleo 2004, Sanchez et al. 2009, Vanlerberghe et al. 2010). All approaches should be tailored to adapt to local conditions depending on the level of urbanization with increasing horizontal management in more rural areas to increase capacity and sustainability (Toaliu and Taleo 2004, Sanchez et al. 2009, Kittayapong et al. 2012, Castro et al. 2012).

##### Community-led surveillance and control of malaria

The literature pulled for malaria indicates that the utilization of community members to conduct both vector surveillance and control activities is proving most effective. Many of these activities involve community health workers, working groups, community education and mobilization, and/or community-led larviciding, indoor residual spraying (IRS), and/or larval source management (LSM) (Johns et al. 2016, Ingabire et al. 2017, Ingabire et al. 2016, Abejirinde et al. 2018, Asale et al. 2019, Chaki et al. 2011), many of which are commonly used within other disease systems as well such as dengue fever. There were benefits and drawbacks seen with many of the CBMs chosen. Community-based IRS was overall found to have easy integration into the current community-based health system in Ethiopia and was equal if not more efficient than the district-based IRS without compromising quality (Johns et al. 2016). While a LSM intervention comparing the effectiveness of community led program integration versus a project research team led program integration saw increases in community willingness to participate in future LSM activities and saw a decrease in mosquito abundance and nuisance biting (Ingabire et al. 2017). An additional study focusing on LSM trained community-members to conduct preliminary mapping of habitats, conduct larvae surveillance, and apply larvicide where larvae were found resulted in a consistent decline in malaria prevalence and mosquito density across the pilot area as the pilot was scaled up to almost 63 districts in Ethiopia (Chaki et al. 2011, 2014). Lastly, the benefits of community working groups and action teams created an increase in the coverage of information, education, health literacy, and more responsible decision making regarding malaria prevention and control due to direct involvement of various members of the community including farmers, youth, military service men, faith leaders, and more (Ingabire et al. 2017,Abejirinde et al. 2018, Asale et al. 2019). Benefits of the above mentioned CBMs revolve around increases in the healthcare workforce capacity, increases in sociocultural understanding, and more strengthened health systems through utilization of the vast, untapped wealth of knowledge within local community members (Ingabire et al. 2017). As has been seen in other disease systems, the main limitations were a lack of formalization of community health workers, participant recruitment, and motivation (Abejirinde et al. 2018, Asale et al. 2019, Sikaala et al. 2014). If informal health workforces, such as community action teams or working groups are being utilized, it is recommended that these informal health teams become part of the formal workforce in order to benefit from government supervision and support. Additionally, increasing the healthcare workforce in this manner will allow for potential expansion of these programs into other health issues such as maternal and child health, nutrition, and WASH (Abejirinde et al. 2018). When focusing on recruitment and motivation, it is recommended that community leaders conduct participant recruitment processes, to recruit participants from existing vector control programs, and to focus participant incentives and motivation through promotions or raises to increase community ownership (Fillinger et al. 2008, Chaki et al. 2011). For malaria surveillance and control overall, a system that mobilizes and involves community members, is integrated into current vertical systems, and managed by either local or municipal health centers is recommended given its engagement of varying levels of stakeholders and potential for sustainability (Chaki et al. 2014). It is important to note that previous approaches of IVM, which simply delivered proven routine interventions through a trained professional, are not sustainable without involvement of the community (Mukabana et al. 2006). Overall, the direct involvement of community action teams or working groups helped sensitize communities to IVM strategies and vector control interventions that are less common in malaria endemic areas of the world, such as environmental management and source reduction (Ingabire et al. 2017). The framing of this literature shows that an IVM strategy can be beneficial to reduce malaria beyond what is already achieved by routine interventions, but the approach must always be implemented in the context of the countries specific vector situation, complementary to current interventions, and should be flexible to change chosen interventions as needed (Ingabire et al. 2016, Johns et al. 2016, Mutero et al. 2020).

##### Citizen Science (CS) focused disease systems Community-based active and passive tick surveillance

Based on the literature on community approaches for tick-borne diseases (TBDs), entomological surveillance through the use of citizen science (CS) initiatives is considered the best way to track the spread of ticks and create robust datasets (Hines and Sibbald 2015, Lewis et al. 2018, Little et al. 2019). One common CS initiative is a form of community submission of ticks, which allows submission to come from either active collection methods such as personal tick dragging in yards or gardens or from passive ones such as removal from self, pets, plants, or livestock (Lewis et al. 2018, Little et al. 2019). The benefits to data collected through CS initiatives is that it can result in high-density, longitudinal data of small areas in which detailed ecological observations of the climate, species, and other driving factors of vector ecology can be better understood and investigated (Lewis et al. 2018, Little et al. 2019, Hart et al. 2022, Porter et al. 2021a). These data do have limitations, however, that revolve mainly around the lack of standardization in collection methods utilized, the potential for inaccurate data because of it, and biases introduced through human behavior and activity (Jongejan et al. 2019, Hart et al. 2022, Foldvari et al. 2022). Another reason to introduce CBMs through CS is to increase community engagement and knowledge. Implementing CS initiatives and increasing community-academic tick surveillance partnerships can help relay scientific knowledge to the public and allow communities to act on this new information (Lewis et al. 2018). It has been shown that partnering with trusted community members can help reduce resistance to public health messaging and can increase awareness of tick-bite prevention practices (Lewis et al. 2018). Another common CS method is the use of mobile phone applications to record tick submissions, the collection site, environmental factors, and more (Hines and Sibbald 2015, Jongejan et al. 2019). These mobile phone applications can also offer information on how to identify ticks and tick habitats, how to protect yourself, basic TBD symptom information, and/or can allow for geotagged photos of ticks to be uploaded and distribution maps to be created (Hines and Sibbald 2015). These applications make available ‘real-time’ information that is easily accessible by the general population, but any mobile phone apps created must be user friendly and advertised to all, not just those interested in vector surveillance and control or public health (Hines and Sibbald 2015). CS is beneficial for increasing the geographic scope of the data collected creating very large, spatially diverse datasets (Porter et al. 2021a, 2021b). The main conclusion found in the review is that areas with limited longitudinal data on ticks and TBD presence, whether they are endemic for TBDs or not, would benefit from implementing a CS initiative and including community members in the data collection process (Lewis et al. 2019, Porter et al. 2021a). It is recommended that future CS projects have clear and simple protocols that allow for consistent participation and motivation from community members and consider the demands of daily life (Lewis et al. 2019, Foldvari et al. 2022, Chenery et al. 2022). Though CS initiatives can be helpful in monitoring pathogen emergence in areas traditionally nonendemic for TBDs, they can also be powerful when combined with conventional surveillance methods in areas with current vector control systems and should be used to supplement current processes to increase sample numbers and monitor the changing dynamics of ticks and TBDs (Jongejan et al. 2019, Porter et al. 2021a, 2021b).

##### General mosquito (exclusive of dengue and malaria) focused citizen science and non-CS initiatives

Based upon the literature in this review, the most common CBM for mosquito-borne diseases, exclusive of dengue and malaria, is conducting surveillance through CS initiatives. Successful CS initiatives seen for MBD include the GLOBE Mosquito Habitat Mapper (Low et al. 2021, Freeman et al. 2022) and the Mozzie Monitors project on the iNaturalist platform (Sousa et al. 2022). These platforms encourage citizens to participate in public health research, to help spread correct information regarding VBDs, and are helpful in creating large, diverse datasets (Tarter et al. 2019, Low et al. 2021, Freeman et al. 2022, Sousa et al. 2022).The Mozzie Monitors project allowed users to upload images of adult mosquitoes and yields information about diversity and species distribution on an already established platform with a strong user base (Sousa et al. 2022). On the other hand, the GLOBE Mosquito Habitat Mapper project allows users to document potential breeding grounds and primarily captures data on larval stages and helps bridge surveillance gaps in *Aedes* distribution in West Africa (Freeman et al. 2022). One specific study conducted by Carney, et.al, was able to integrate the datasets from separate citizen science projects into a single dashboard called The Global Mosquito Observations Dashboard that can be used to target and track all mosquito species, including invasive ones, all while testing and using artificial intelligence (AI) systems to help identify mosquito species (Carney et al. 2022). The datasets combined in the dashboard include Mosquito Alert (Bartumeus et al. 2018), iNaturalist (Cull 2021, Sousa et al. 2022), and the GLOBE observer (Low et al. 2021, Freeman, et al. 2022), all of which can be seen in other papers throughout this scoping review. The Global Mosquito Observations Dashboard serves as a way for citizens and professionals to collaborate in the fight against mosquito-borne diseases worldwide. The dashboard can be accessed at mosquitodashboard.org (GMOD (arcgis.com)). Focus for CS initiatives needs to be on engaging users, continuing to gather new users across all socio-economic statuses, and on photo submission and identification protocol (Low et al. 2021, Freeman et al. 2022, Sousa et al. 2022). These methods are increasing in popularity as the use of technology increases across the world with improvements in internet access, smartphones, GPS, and high-resolution cameras (Sousa et al. 2020, Cull 2021). These platforms and dashboards are not limited by geographic boundaries or time unlike conventional active surveillance methods and can still gather information on temporality and seasonality in an economic and more epidemiologically relevant way (Babu et al. 2019, Little et al. 2019). But there are limitations revolving mainly around motivation and participation, potential spatial bias to where humans frequent, technological issues such as camera quality, internet access, user accessibility on the app, and potential start-up costs (Curtis-Robles et al. 2015, Murindahabi et al. 2018, Little et al. 2019, Murindahabi et al. 2021, Chenery et al. 2022). An additional common CBM for MBD surveillance outside of CS seen in the literature includes mosquito trapping conducted through CS methods where community members are mailed or given mosquito traps such as the BG GAT trap (Sousa et al. 2020) or the BG Sentinel trap (Craig et al. 2021). Householders were trained on how to set up and conduct surveillance with the trap within their household setting and sent results back to researchers. The BG GAT trap was sent to Australian residents and was considered effective in mosquito collection, easy to ship, required little to no power, and cost less than a trained entomologist (Sousa et al. 2020). The BG sentinel trap in the Solomon Islands found mosquito identification by community members to have high levels of agreement to a trained entomologists assessment suggesting that community members can identify mosquitoes with an adequate level of accuracy (Craig et al. 2021).

Success of these types of methods relies on detailed consideration of access to electricity, access to technology such as a mobile phone to send or upload results, and ease of use (Sousa et al. 2020, Craig et al. 2021). There have also been numerous CS initiatives focused on nuisance biting mosquitoes in Europe like the Mosquito Reporting Scheme in the UK, Muckenatlas in Germany, Muggenradar in The Netherlands, AtrapaelTigre.com in Spain, iMoustique in France, and MosquitoWEB in Portugal (Kampen et al. 2015). These initiatives have all been beneficial in gathering up-to date occurrence and distribution data in Europe.

## Discussion

Over time, CBMs have been seen as an engaging way to increase acceptability and sustainability of vector surveillance and control interventions through direct community involvement. This scoping review describes the current landscape of CBMs and its successes within various disease systems.

Notable successes include increases in surveillance coverage and the creation of large, diverse datasets not possible through traditional surveillance alone (Rojas-Cortez et al. 2016, Tarter et al. 2019, Abrahan et al. 2021, Low et al. 2021, Freeman et al. 2022, Sousa et al. 2022,), increased community knowledge, engagement, ownership, & sustainability (Ingabire et al. 2017, Lewis et al. 2018, Asale et al. 2019, Abrahan et al. 2021), and lastly, increases in the healthcare workforce & a more strengthened health system (Ingabire et al. 2017, Asale et al. 2019, Abrahan et al. 2021). But challenges are still present and across disease systems the major challenges revolve around individual participation and motivation (Hashimoto and Yoshioka 2012, Lewis et al. 2019, Sanchez et al. 2009, Abejirinde et al. 2018).

A big distinction made in the results section is between citizen science (CS) and CBMs. CS and CBMs often get mistaken as interchangeable terms; however, while all citizen science initiatives utilize the community, they are not foundationally the same as CBMs. The main foundation that differentiates CS from other forms of CBMs or community engagement is that participants voluntarily search for ways to help conduct scientific research. It is true that participants of CBMs / community engagement agree to participate and do so willingly, they most often were advertised too and approached to participate versus seeking out the program or intervention of their own personal interest. An additional difference is that CS is most often ad hoc with no clear health system structure while CBMs coordinate centrally with clear time points. On the other hand, these two methods have many aspects in common such as involving members of the general public or non-professionals in data collection with the focus of improving scientific research and public knowledge.

Of the chosen disease systems, both Chagas disease and malaria made up the largest groups of literature published in this review with chagas disease predominately in Central and South America and malaria in Africa. A focus of this review being on the defined publication categories revealed that the ‘community participation’ category was the most common with direct utilization with the chagas disease field. Throughout the results section, the important distinction between vector control methods and vector surveillance methods is made clear within each disease system noting that surveillance methods are predominately utilized in chagas disease, TBDs, and MBDs (exclusive of dengue and malaria), while vector control methods are predominately utilized in the dengue field, leaving the utilization of both vector control and surveillance within the field of malaria. The dual approach within malaria is focused primarily on Larval Source Management (LSM) techniques, whose foundation in environmental management is a common technique in other disease systems such as dengue. Though LSM is not a commonly recommended or supported approach, it has numerous benefits that have been seen in the dengue field and are being seen in research in the malaria field including within the successful community-based program in Tanzania, the Urban Malaria Control Program (UMCP).

The UMCP was a program that was community-based and run by community-owned resource persons but was vertically managed. The program saw success my integrating fully into the decentralized administrative system in Dar es Salaam and through coordination by the City Medical Office of Health which gave the city council ownership over the program (Chaki et al. 2011). Similar to many of the studies presented in this research across all the disease systems, the UMCP faced challenges regarding participant motivation, training, and retention and placed large programmatic focus on community ownership, the recruitment processes, government involvement & integration, geographic & social boundaries, and sustainability (Fillinger et al. 2008, Chaki et al. 2011, 2014). The UMCP serves as a great case-study of a community-based program which involves both surveillance and control and required high levels of community engagement and ownership to be effective as well as being some of the only literature to describe and acknowledge their choice to compensate the participants and the benefits and drawbacks of doing so. Additionally, though not a main reason why the UMCP is highlighted here, the focus on interventions targeting larval stages instead of adult stages is important in the context of an IVM strategy and should be considered as the entomological landscape changes. To read more detailed information about the UMCP and the challenges they faced and overcame, see supplemental document 2.

The current approach to reducing VBD is one that focuses on improving and implementing vector surveillance and control interventions for individual disease systems instead of interventions that can be effective across disease systems. By focusing on individual systems, the full potential of vector surveillance and control programs is not being reached. The individualistic approach often results in poor utilization of resources, a lack of full integration into health systems, is not as adaptable to changing environments, lacks collaboration with other sectors, and can often have a small collection of interventions to choose from (WHO 2012). By taking an integrated vector management (IVM) approach, vector surveillance and control can be repositioned as a key approach to reducing VBD that is cost-effective and sustainable (WHO 2017). The key elements of IVM are the promotion of the use of a wide range of interventions, understanding the importance of local knowledge, addressing several diseases at the same time due to the effectiveness of some interventions against several vectors, and encouraging collaboration within and outside of the health sector (WHO 2012). Unfortunately, since its introduction in 2008, the IVM strategy has received little support due to a lack of political and financial will to reorient current vector surveillance and control programs (WHO 2017). A recent example of the importance of IVM is the introduction of invasive mosquito, *Anopheles stephensi*, into Africa in 2012.

*Anopheles stephensi* has been shown to be resistant to most of the current insecticides used against adult mosquitoes, it is a competent vector for both *Plasmodium vivax* and *Plasmodium falciparum*, and unlike most other *Anopheles* mosquitoes, it can thrive in artificial containers in urban environments similar to *Aedes* mosquitoes (PMI 2023). The response against *An. stephensi* has been slow due to a required pivot in surveillance methods from adult to larval surveillance, which is not often employed for other African malaria vectors. Conversations surround whether or not an IVM approach, one that already employs the infrastructure to implement common surveillance and control methods for *Aedes* mosquitoes towards *Anopheles* mosquitoes, would have developed a quicker response to the spread of *An. stephensi* (PMI 2023). Due to the evolving situation, this conversation should remain high priority.

### Limitations

As with most studies, the design of this scoping review is subject to limitations. First, the publications chosen are limited to arthropod based diseases and those publications written in English. This excludes any publications focusing on non-arthropods vectors such as aquatic snails and fleas.

Additionally, there is potential that the chosen methodology and key words such as “vector control” and “community-based control” utilized to pull the literature overlooked other types of community-based methods of surveillance and control such as those focused on habitat remediation. To increase publications, future studies should consider broadening the scope or focusing on a specific disease and including more inclusive key words. A second limitation is that this review does not focus on the quality of the data or conduct meta-analyses given the broad scope of the data collected and its inability to be systematically analyzed. To improve upon that, future research should consider conducting meta-analyses by narrowing the scope of the review. Lastly, there were a lack of studies reporting that CBMs did not work. This could suggest potential for publication bias, or a lack of studies objectively examining pros and cons of CBMs on this topic.

## Conclusion

Overall, the majority of publications in the review presented positive operational benefits of using community-based methods. As it becomes more common to utilize community members in research in any capacity, it will be critical for researchers to acknowledge the added benefit or drawback that including the community had on their intended outcome or data quality. Doing so will help fill in the gaps that were brought forth in this review, further helping create the public health case for CBMs as a vector surveillance and control tool. Taken together, this scoping review highlights potential challenges and benefits of decentralized community based and citizen science entomological surveillance and control programs and outlines potential future directions for incorporating CBMs into VBD control and elimination programming, and potential for community based IVM approaches.

## Disclaimer

The findings and conclusions expressed herein are those of the authors and do not necessarily represent the official position of the U.S. Centers for Disease Control and Prevention (CDC), U.S. Agency for International Development (USAID), or U.S. President’s Malaria Initiative (PMI). Financial support for PE, SZ, MY, and TA was provided by PMI.

## Supplemental Information

**Supplemental Figure 1:**
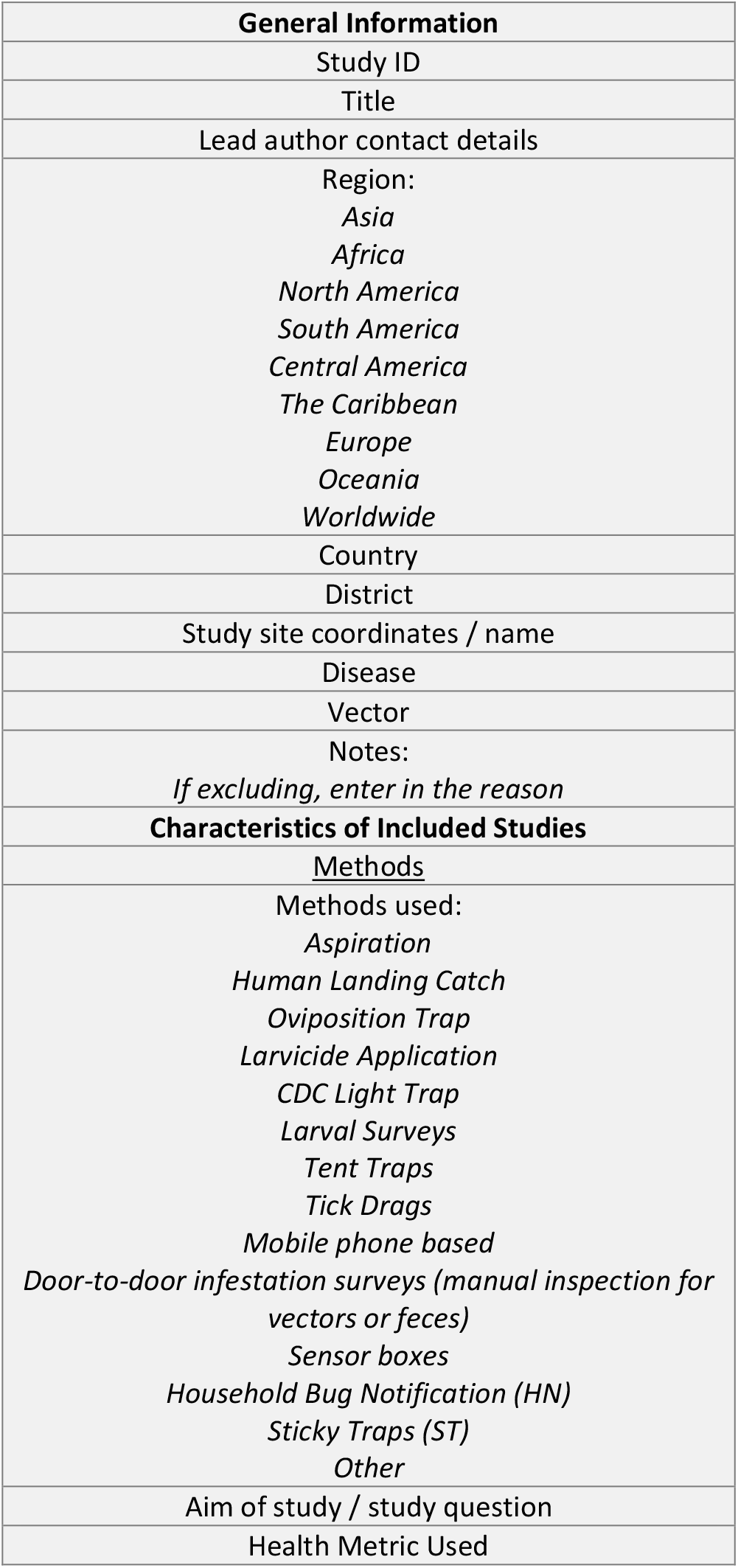
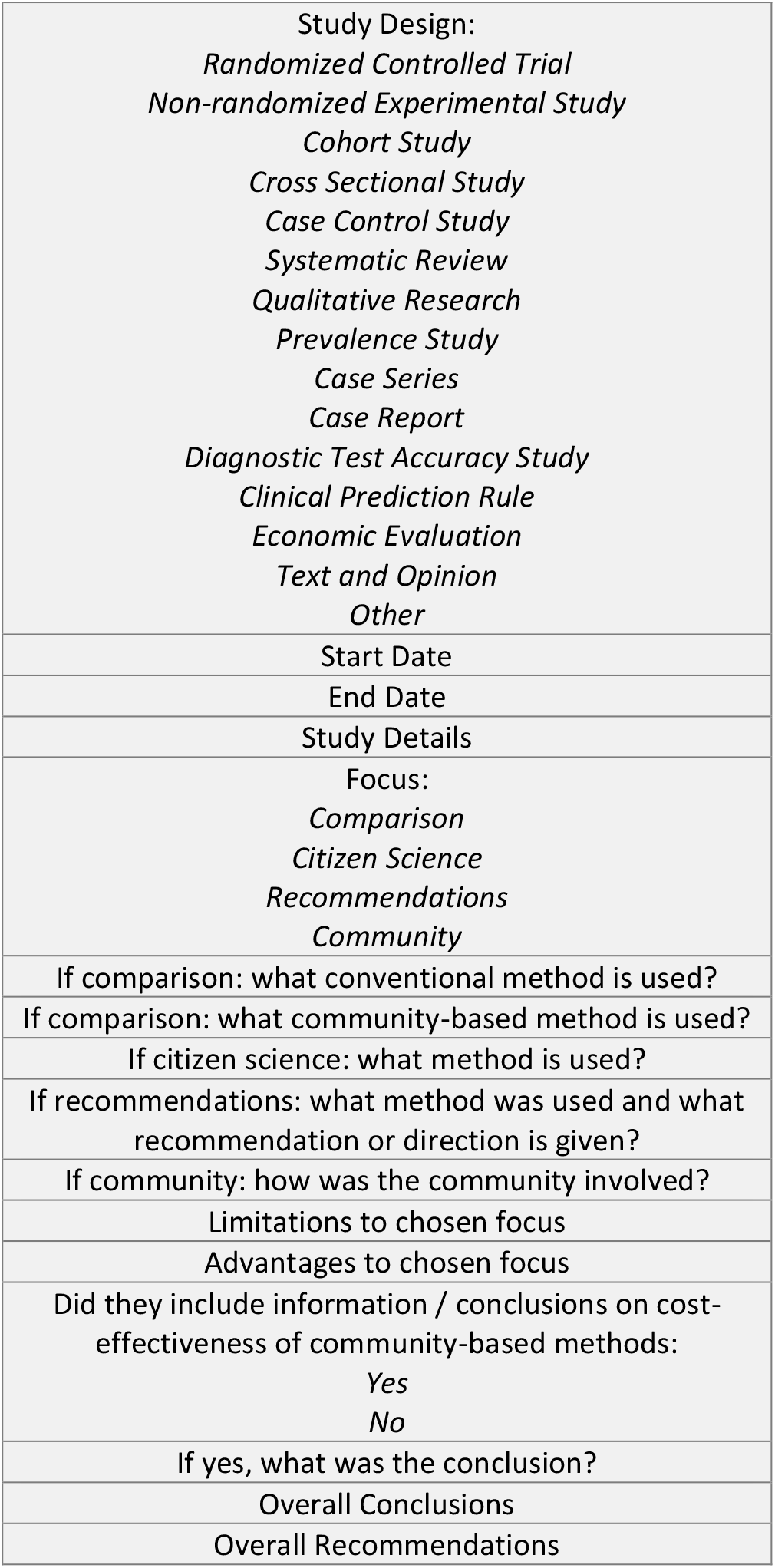
Data Extraction Template

**Supplemental Table 1:**
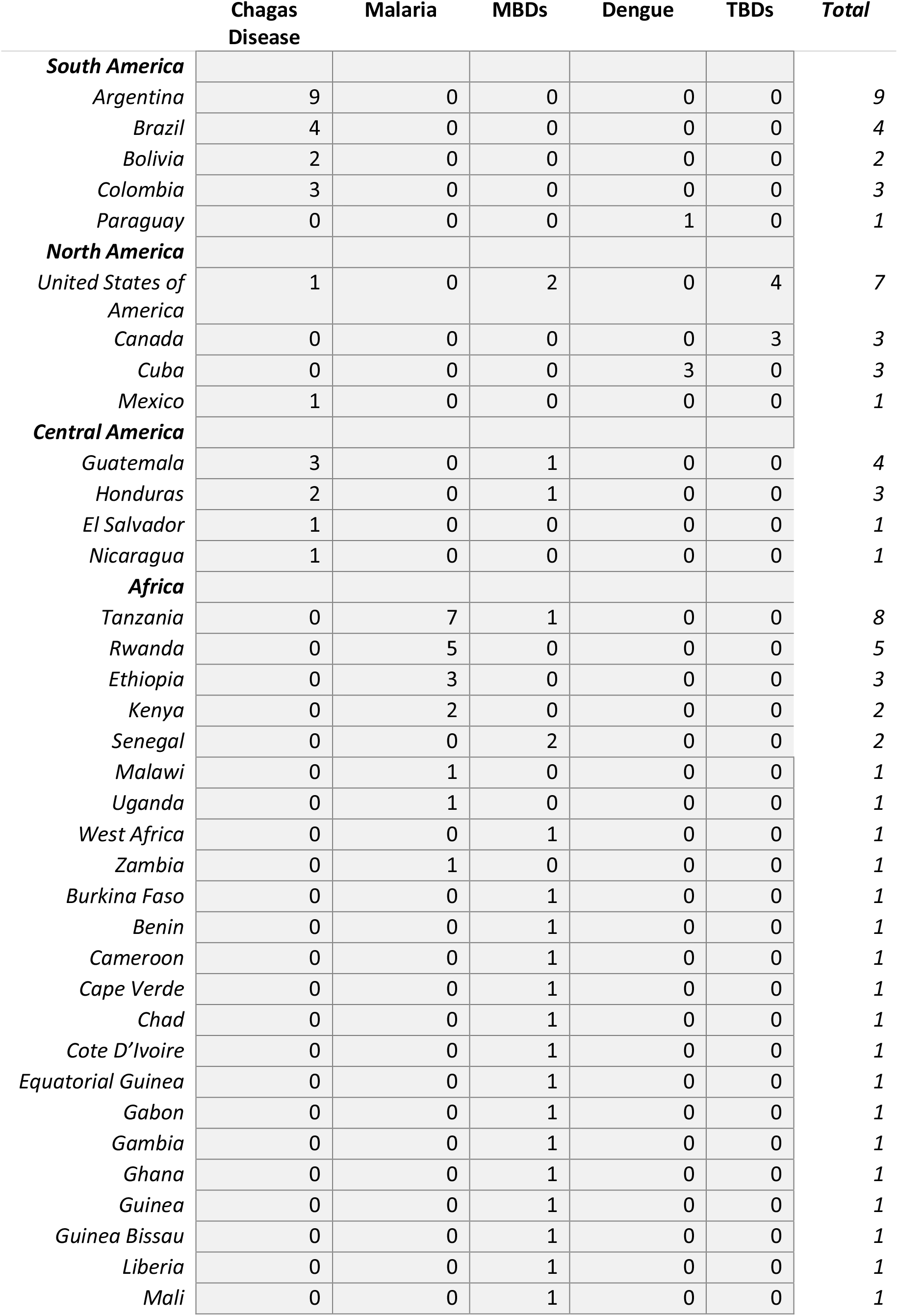
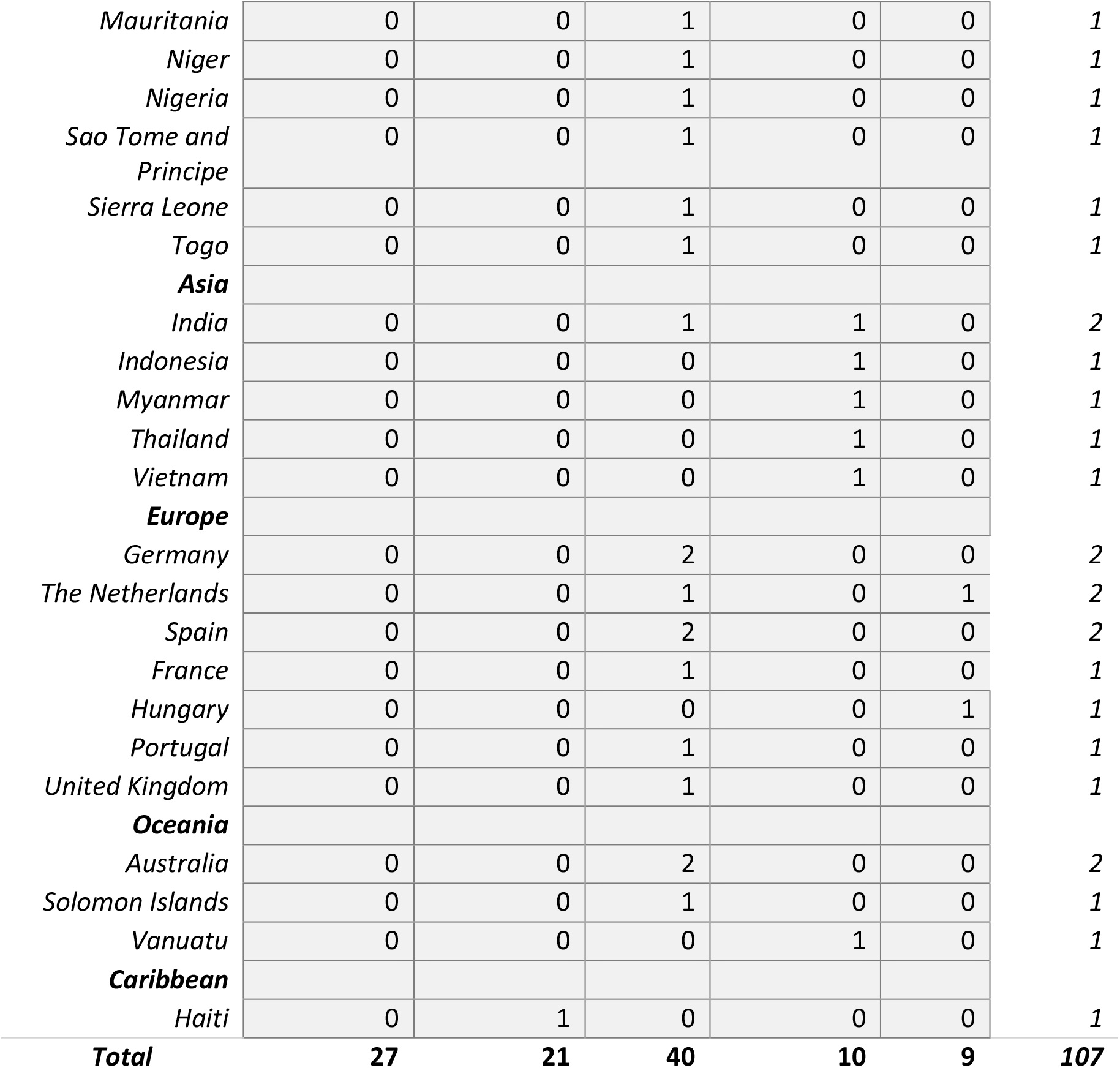
Studies included in final analysis categorized by region of the world and country (n=107) *some studies are conducted in more than one country

|  | Chagas<br>Disease | Malaria | MBDs | Dengue | TBDs | Total |
| --- | --- | --- | --- | --- | --- | --- |
| **South America** |  |  |  |  |  |  |
| Argentina | 9 | 0 | 0 | 0 | 0 | 9 |
| Brazil | 4 | 0 | 0 | 0 | 0 | 4 |
| Bolivia | 2 | 0 | 0 | 0 | 0 | 2 |
| Colombia | 3 | 0 | 0 | 0 | 0 | 3 |
| Paraguay | 0 | 0 | 0 | 1 | 0 | 1 |
| **North America** |  |  |  |  |  |  |
| United States of<br>America | 1 | 0 | 2 | 0 | 4 | 7 |
| Canada | 0 | 0 | 0 | 0 | 3 | 3 |
| Cuba | 0 | 0 | 0 | 3 | 0 | 3 |
| Mexico | 1 | 0 | 0 | 0 | 0 | 1 |
| **Central America** |  |  |  |  |  |  |
| Guatemala | 3 | 0 | 1 | 0 | 0 | 4 |
| Honduras | 2 | 0 | 1 | 0 | 0 | 3 |
| El Salvador | 1 | 0 | 0 | 0 | 0 | 1 |
| Nicaragua | 1 | 0 | 0 | 0 | 0 | 1 |
| **Africa** |  |  |  |  |  |  |
| Tanzania | 0 | 7 | 1 | 0 | 0 | 8 |
| Rwanda | 0 | 5 | 0 | 0 | 0 | 5 |
| Ethiopia | 0 | 3 | 0 | 0 | 0 | 3 |
| Kenya | 0 | 2 | 0 | 0 | 0 | 2 |
| Senegal | 0 | 0 | 2 | 0 | 0 | 2 |
| Malawi | 0 | 1 | 0 | 0 | 0 | 1 |
| Uganda | 0 | 1 | 0 | 0 | 0 | 1 |
| West Africa | 0 | 0 | 1 | 0 | 0 | 1 |
| Zambia | 0 | 1 | 0 | 0 | 0 | 1 |
| Burkina Faso | 0 | 0 | 1 | 0 | 0 | 1 |
| Benin | 0 | 0 | 1 | 0 | 0 | 1 |
| Cameroon | 0 | 0 | 1 | 0 | 0 | 1 |
| Cape Verde | 0 | 0 | 1 | 0 | 0 | 1 |
| Chad | 0 | 0 | 1 | 0 | 0 | 1 |
| Cote D'Ivoire | 0 | 0 | 1 | 0 | 0 | 1 |
| Equatorial Guinea | 0 | 0 | 1 | 0 | 0 | 1 |
| Gabon | 0 | 0 | 1 | 0 | 0 | 1 |
| Gambia | 0 | 0 | 1 | 0 | 0 | 1 |
| Ghana | 0 | 0 | 1 | 0 | 0 | 1 |
| Guinea | 0 | 0 | 1 | 0 | 0 | 1 |
| Guinea Bissau | 0 | 0 | 1 | 0 | 0 | 1 |
| Liberia | 0 | 0 | 1 | 0 | 0 | 1 |
| Mali | 0 | 0 | 1 | 0 | 0 | 1 |
| *Mauritania* | 0 | 0 | 1 | 0 | 0 | 1 |
| *Niger* | 0 | 0 | 1 | 0 | 0 | 1 |
| *Nigeria* | 0 | 0 | 1 | 0 | 0 | 1 |
| *Sao Tome and Principe* | 0 | 0 | 1 | 0 | 0 | 1 |
| *Sierra Leone* | 0 | 0 | 1 | 0 | 0 | 1 |
| *Togo* | 0 | 0 | 1 | 0 | 0 | 1 |
| **Asia** |  |  |  |  |  |  |
| *India* | 0 | 0 | 1 | 1 | 0 | 2 |
| *Indonesia* | 0 | 0 | 0 | 1 | 0 | 1 |
| *Myanmar* | 0 | 0 | 0 | 1 | 0 | 1 |
| *Thailand* | 0 | 0 | 0 | 1 | 0 | 1 |
| *Vietnam* | 0 | 0 | 0 | 1 | 0 | 1 |
| **Europe** |  |  |  |  |  |  |
| *Germany* | 0 | 0 | 2 | 0 | 0 | 2 |
| *The Netherlands* | 0 | 0 | 1 | 0 | 1 | 2 |
| *Spain* | 0 | 0 | 2 | 0 | 0 | 2 |
| *France* | 0 | 0 | 1 | 0 | 0 | 1 |
| *Hungary* | 0 | 0 | 0 | 0 | 1 | 1 |
| *Portugal* | 0 | 0 | 1 | 0 | 0 | 1 |
| *United Kingdom* | 0 | 0 | 1 | 0 | 0 | 1 |
| **Oceania** |  |  |  |  |  |  |
| *Australia* | 0 | 0 | 2 | 0 | 0 | 2 |
| *Solomon Islands* | 0 | 0 | 1 | 0 | 0 | 1 |
| *Vanuatu* | 0 | 0 | 0 | 1 | 0 | 1 |
| **Caribbean** |  |  |  |  |  |  |
| *Haiti* | 0 | 1 | 0 | 0 | 0 | 1 |
| **Total** | **27** | **21** | **40** | **10** | **9** | **107** |

**Supplemental Table 2:**
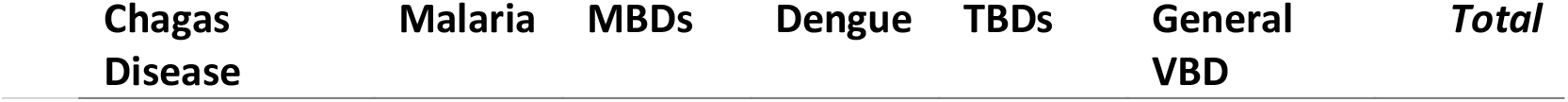
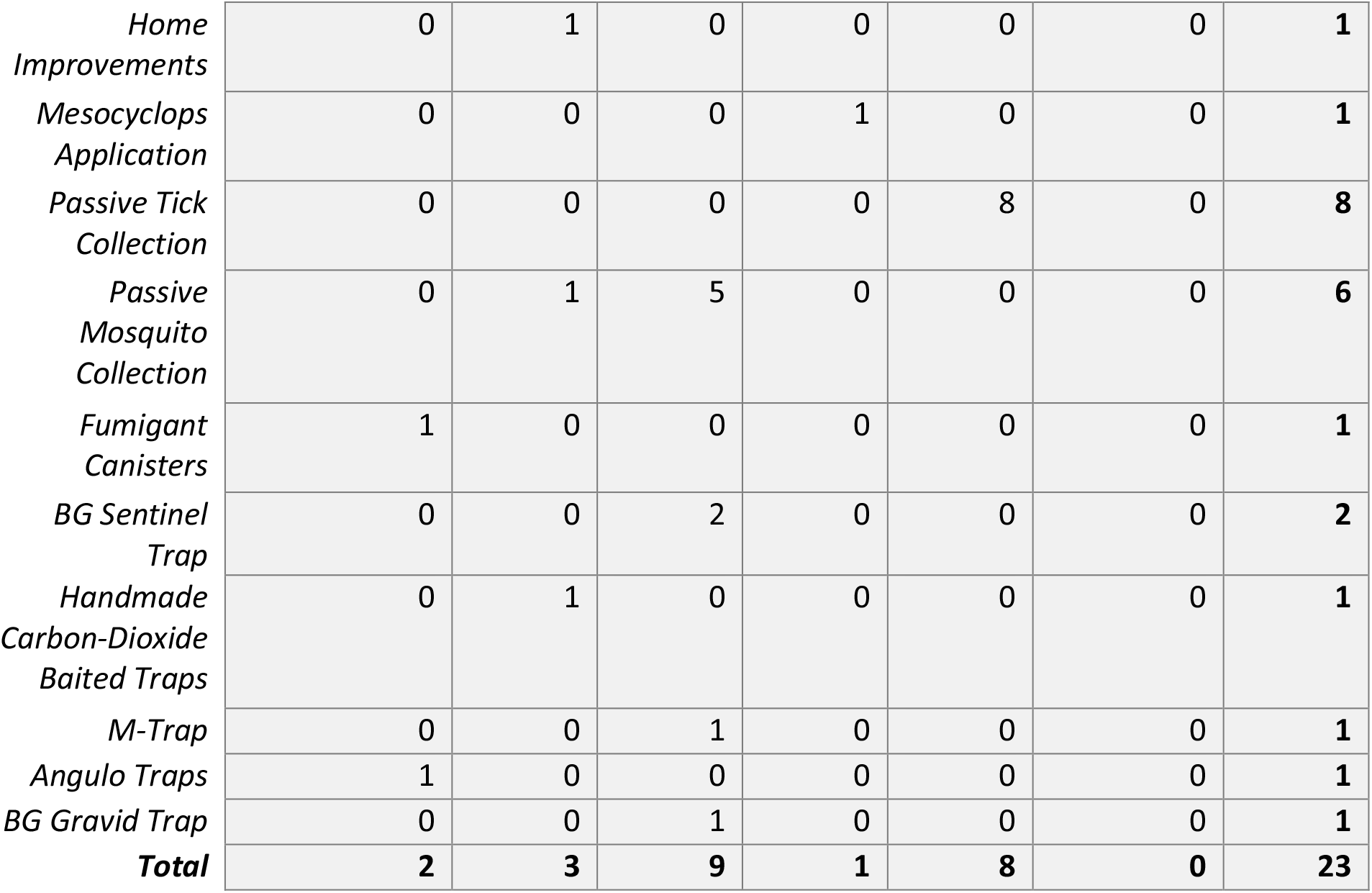
Breakdown of Capture Method - “Other” by Disease

| Chagas Disease | Malaria | MBDs | Dengue | TBDs | General VBD | Total |
| --- | --- | --- | --- | --- | --- | --- |

|  |  |  |  |  |  |  |  |
| --- | --- | --- | --- | --- | --- | --- | --- |
| *Home Improvements* | 0 | 1 | 0 | 0 | 0 | 0 | **1** |
| *Mesocyclops Application* | 0 | 0 | 0 | 1 | 0 | 0 | **1** |
| *Passive Tick Collection* | 0 | 0 | 0 | 0 | 8 | 0 | **8** |
| *Passive Mosquito Collection* | 0 | 1 | 5 | 0 | 0 | 0 | **6** |
| *Fumigant Canisters* | 1 | 0 | 0 | 0 | 0 | 0 | **1** |
| *BG Sentinel Trap* | 0 | 0 | 2 | 0 | 0 | 0 | **2** |
| *Handmade Carbon-Dioxide Baited Traps* | 0 | 1 | 0 | 0 | 0 | 0 | **1** |
| *M-Trap* | 0 | 0 | 1 | 0 | 0 | 0 | **1** |
| *Angulo Traps* | 1 | 0 | 0 | 0 | 0 | 0 | **1** |
| *BG Gravid Trap* | 0 | 0 | 1 | 0 | 0 | 0 | **1** |
| **Total** | **2** | **3** | **9** | **1** | **8** | **0** | **23** |

