## Supplemental document 1 for "Community-Based Entomological Surveillance and Control of Vector-Borne Diseases: A Scoping Review"

### *Determine your starting point*

**DIRECTIONS:** Below, split by disease, are the main recommendations / directions that the literature has presented including references to support them.

**CONTENT:** Each disease has a section included in this document to cover the main recommendation, key points / items to consider, potential challenges, benefits, and optimal environment

| Disease | Main Direction | References | |
|---|---|---|---|
| **MALARIA** SECTION A | Main Direction: Compensated community members conducting surveillance and control through larval source management | Chaki, 2011<br>Chaki, 2009<br>Fillinger, 2008<br>Vanek, 2006 | Geissbuhler, 2009<br>Chaki, 2014<br>Dongus, 2007<br>Mukabana, 2006 |
| **DENGUE FEVER** SECTION B | Main Direction: Community empowerment and education on source reduction techniques taught by community members | Castro, 2012<br>Tana, 2012<br>Nam, 2004<br>Sommerfeld, 2015 | Toaliu, 2004<br>Kittayapong, 2012<br>Sanchez, 2009<br>Vanlerberghe, 2010<br>Wai, 2012 |
| **CHAGAS DISEASE** SECTION C | Main Direction: Householder notification and supplementary use of various detection devices - sensor boxes, sticky tape, etc. | Abrahan, 2021<br>Cecere, 2019<br>Enriquez, 2020<br>Hashimoto, 2015<br>Leon, 2019 | Weeks, 2014<br>Abad-Franch, 2011<br>Cardinal, 2007<br>Carvallo, 2018<br>Dumonteil, 2019 |
| **TICK-BORNE DISEASES** SECTION D | Main Direction: Citizen Science initiatives focused on active & passive surveillance utilizing technology such as mobile phones | Egizi, 2019<br>Hart, 2020<br>Lewis, 2018<br>Porter, 2021<br>Hines, 2015 | Chenery, 2022<br>Little, 2019<br>TannerPorter, 2021<br>Jongejan, 2019<br>Foldvari, 2022 |
| **MOSQUITO-BORNE DISEASES*** SECTION E | Main Direction: Citizen Science initiatives focused on community submission of mosquito samples or photos utilizing either mosquito trapping devices or mobile phone applications | Craig, 2021<br>Werner, 2020<br>Low, 2021<br>Freeman, 2022<br>Sousa, 2022 | Carney, 2022<br>Bartumeus, 2018<br>Cull, 2021<br>BrazSousa, 2020<br>Kampen, 2015 |

*Mosquito-borne diseases exclusive of malaria and dengue

### Section A: Malaria

**MAIN RECOMMENDATION** A vector surveillance and control program that is integrated into a current vertical system that selects, trains, and compensates local community members in conducting larval source management (including larval surveys and larviciding) in their respective wards or neighborhoods and that is managed by either local or municipal health centers

| **BACKGROUND:** | Larval source management (LSM) has historically been a gold standard and routine part of Aedes mosquito control. But as a method for Anopheles mosquito control, it has a limited scope per the WHO recommendations and is currently considered supplementary to routine methods. As urban human population in Africa continues to grow and cases of malaria increase, LSM should be considered again as an option for anopheles mosquito control due to its ability to be implemented easily in urban environments and to target a larger portion of the mosquito population. In order for LSM vector control methods such as larviciding to be successful, rigorous vector surveillance of aquatic stage mosquitoes is needed. Involving the community in the LSM process taps into a large reservoir of untapped knowledge, spreads knowledge about malaria within communities, and can potentially be low cost.<br>These community members would be effectively chosen from the local area by community leaders and groups, adequately trained and supervised, and responsible for conducting various types of surveillance and larval source management techniques. It is worth noting that the above recommendations can also be applied to other anopheles-based diseases such as lymphatic filariasis but must still be considered within its own context.<br><br>**Best example: The Urban Malaria Control Program in Dar es Salaam, Tanzania (1-8)** |
|---|---|

**Key points / items to consider:**
Community-wide acceptance & ownership
Recruitment of participants by local leadership within intervention areas
Compensation comparable to workload
Quality of training
Supervision/management quality and workload

**Challenges that might arise:**
Access to private areas / fenced areas
Lack of communication skills training
Participant retention / satisfaction

**Benefits of chosen method:**
Complete utilization of local knowledge
Potentially cost-effective
Project sustainability
Incorporates IVM strategy

**Optimal Environment:**
Areas with focal and low to moderate transmission
Urban areas / areas with an existing administrative boundary system

### Section B: Dengue Fever

**MAIN RECOMMENDATION** A blended program involving current biological or chemical larval control tools at the vertical level and a horizontal program involving household managed source reduction and a community empowerment program

| BACKGROUND: | Historically, dengue control has been viewed as the government's responsibility due to the specific focus on technical vector control interventions that do not involve the community such as mass larviciding campaigns or large-scale environmental management. But the combination of aedes mosquitoes' preference to breed in any and all containers and an increase in urbanization, has made it hard for the government and local municipalities to control the vector without community involvement. In order to increase local knowledge about dengue and potentially create behavior changes, environmental management techniques such as source reduction combined with community empowerment strategies are highly recommended due to the fact that they can be completed by anyone consistently, with minimal training, and encourages capacity building.<br><br>Allowing community members to create working groups who are responsible for providing education on source reduction either through door-to-door or community workshops has proven to be effective ways to reduce larval levels of container breeding mosquitoes. It is worth noting that the below recommendations can also be applied to other aedes-based diseases such as zika virus, yellow fever, and chikungunya but must still be considered within their own contexts.<br><br>**Best example: Dengue vector control and community empowerment projects in Cuba (9, 15, 16)** |
|---|---|

**Key points / items to consider:**

Recruitment for working groups
Training quality
Quality educational materials

**Challenges that might arise:**

Community member motivation to continue source reduction
Collaboration with other municipal authorities such as waste and sanitation
Continued support from local government

**Benefits of chosen method:**

Increased community knowledge
Behavior changes
Potentially cleaner communities due to elimination of unused containers

**Optimal Environment:**

Anywhere with interested community members
Urban Environments

### Section C: Chagas Disease

**MAIN RECOMMENDATION** A combination of surveillance methods, such as householder notification and detection devices (sticky tape, sensor boxes, etc.) resulting in a targeted vertical attack phase, such as an insecticide spraying campaign

**BACKGROUND:**

The goal of chagas disease elimination in humans revolves around human dwellings and surrounding structures due to triatomines preference to live in the walls and cracks of structures. Transmission of chagas disease is mainly prevented through reducing house infestation and re-infestation of triatomines and insecticide spraying of specimen positive homes. This is best achieved through a continual surveillance system in the home that even runs during the nighttime when triatomines are most active and has a high sensitivity and effectiveness for accurate detection of infestations. Research has shown that both householder notification of triatomines and placement of detection devices can be as or more sensitive than the gold-standard of timed-manual collection by a trained professional and can be implemented consistently, with minimal training, and at a lower cost. Utilizing both methods allows for detection of both active and resting triatomines as well as the identification of defecation or urination which indicates potential infestation. Additionally, householder notification and detection devices help elimination programs target insecticide spraying to only the necessary homes.

Householder notification and the placement of detection devices involves community members conducting both active and passive surveillance in their homes and collecting and delivering any suspected triatomine bugs to local health authorities to inform elimination programming.

**Best example: Comparison of householder notification, sensor boxes, and timed-manual searches in Central & South America (20, 23)**

**Key points / items to consider:**

Training quality
Materials given to collect triatomines
Specimens drop off / collection points
Domestic versus peri-domestic spaces

**Challenges that might arise:**

Ensuring continued motivation
Quick responses to positive infestations by national programs
Installation and use of detection devices

**Benefits of chosen method:**

Increased coverage
More accurate insecticide spraying campaigns
Reduced re-infestations

**Optimal environment:**

Areas with low-infestation rates
Rural / remote areas

### Section D: Tick-Borne Diseases

**MAIN RECOMMENDATION** Citizen Science initiatives involving active & passive collection of ticks by community members often in combination with a mobile phone application to gather geographic and morphological data

| BACKGROUND: | The transmission cycle for tick-borne diseases is complex and includes both the vector, a reservoir host, and a human host. Due to this complex transmission cycle, an increase in the frequency of ticks, and their expanding geographic reach due to climate change - very robust and detailed data are needed to draw conclusions and make recommendations for protection. The current gold standard for tick surveillance, the tick drag, requires a skilled technician, only targets specific areas at specific times, and can be costly. Citizen science initiatives that encourage passive and active surveillance from the general public provide large datasets that are created continuously, rapidly, and are made of spatially diverse samples. These datasets can be used to gather important information on presence, abundance, and distribution of invasive species and in areas where ticks might be emerging.<br><br>One of the most common methods utilized in these initiatives are mobile phone apps that can identify and geolocate specimens to create distribution maps and more. These apps and their associated data can be critical to areas where infrastructure and funding are lacking for local officials to conduct surveillance as well as areas with developed routine surveillance systems to supplement them. Citizen science is recommended for TBD research because it helps cover large geographic areas, is a consistent form of surveillance, costs less, and increases public knowledge and trust in vector control programs. These<br><br>**Best example: Passive & active citizen science methods in Europe (37) & USA (31)** |
|---|---|

**Key points / items to consider:**
Clear and precise protocol
Consider local access to technology
Advertising of opportunity

**Challenges that might arise:**
Ensuring continued motivation
Lack of standardized collection method
Access to technology / internet access

**Benefits of chosen method:**
Increased coverage
Creation of larger, more diverse datasets
Easier to deploy
Increase in local knowledge

**Optimal environment:**
Areas with limited longitudinal data
Areas without a current vector surveillance infrastructure

### Section E: Mosquito-Borne Diseases*

*Mosquito-borne diseases exclusive of malaria and dengue

**MAIN RECOMMENDATION** Citizen science initiatives that involve community submission of mosquito samples or photos gathered through either a mosquito trapping device or mobile phone application

| BACKGROUND: | |
|---|---|
| | Similar to Tick-Borne Diseases, climate change is increasing the geographic spread of various mosquito species and in conjunction with new invasive species, achieving adequate surveillance is a challenge. In order to either find proof or predict where existing or new invasive mosquito species might spread to, geographically large and entomologically diverse datasets are needed. Based on the research, this is most easily obtained through citizen science initiatives with two common directions of mobile phone applications and mosquito trapping devices. Mobile phone applications are most often used by community members to upload images of larval stage mosquitoes, adult mosquitoes, and potential mosquito breeding grounds. The most common mosquito trapping devices utilized are the BG GAT trap and the BG Sentinel trap. These projects had community members set up, maintain the traps, and upload all gathered samples through mobile phone pictures.<br><br>Citizen science initiatives encourage citizens to participate in public health research, help spread accurate information about VBDs, and are helpful in creating large, diverse datasets. These initiatives can also help gather information on species distribution and diversity that can help bridge surveillance gaps and solve outbreaks before they arise.<br><br>**Best example: Successful initiatives such as the Global Mosquito Observations Dashboard (43) and the BG Sentinel Trapping Scheme in the Solomon Islands (38)** |

**Key points / items to consider:**

Clear and precise photo submission protocol
Consider local access to technology and electricity
Advertising of opportunity
Ease of use of chosen trap

**Challenges that might arise:**

Ensuring continued motivation
Lack of standardized collection method
Access to technology / internet access

**Benefits of chosen method:**

Increased coverage
Creation of larger, more diverse datasets
Increase in local knowledge
Beneficial for studying nuisance mosquitoes

**Optimal environment:**

Areas with limited longitudinal data or limited data in general

**35. Tanner Porter, W., Z. A. Barrand, J. Wachara, K. DaVall, J. R. Mihaljevic, T. Pearson, D. J. Salkeld, and N. C. Nieto. 2021.** Predicting the current and future distribution of the western black-legged tick, Ixodes pacificus, across the Western US using citizen science collections. PLoS ONE 16.

**36.Jongejan, F., S. de Jong, T. Voskuilen, L. van den Heuvel, R. Bouman, H. Heesen, C. Ijzermans, and L. Berger. 2019**. "Tekenscanner": a novel smartphone application for companion animal owners and veterinarians to engage in tick and tick-borne pathogen surveillance in the Netherlands. Parasit Vectors 12: 116.

**37.Foldvari, G., E. Szabo, G. E. Toth, Z. Lanszki, B. Zana, Z. Varga, and G. Kemenesi. 2022.** Emergence of Hyalomma marginatum and Hyalomma rufipes adults revealed by citizen science tick monitoring in Hungary. Transbound Emerg Dis 18: 18.
