## supplemental document 2 for "Community-Based Entomological Surveillance and Control of Vector-Borne Diseases: A Scoping Review"

**Supplemental Document 2: Case study of the Urban Malaria Control Program (UMCP)**

A case study of the Urban Malaria Control Program (UMCP) with larval source management (LSM) in Dar es Salaam, Tanzania serves as the best example of a community-based program implemented in this review. The UMCP involves both surveillance and control and needed high levels of community engagement and ownership to be effective. Though not a main reason why the UMCP was chosen as a case study, the focus on interventions targeting larval instead of adult stages is important in the context of IVM strategy and should continue to be considered as the entomological landscape changes. These changes can be seen with the current rapidly spreading invasive vector *Anopheles stephensi* in Africa, which is well adapted to urban environments, and predominantly occupying man-made breeding habitats (PMI, 2023).

*Community-Based Vector Surveillance and Control Case Study: The Urban Malaria Control Program, Tanzania*

The key management structure of the UMCP, which was one that was community-based but vertically managed, is what made it so successful. The program was fully integrated into the decentralized administrative system and coordinated by the City Medical Office of Health giving the city council ownership over the program (Chaki et al. 2011).

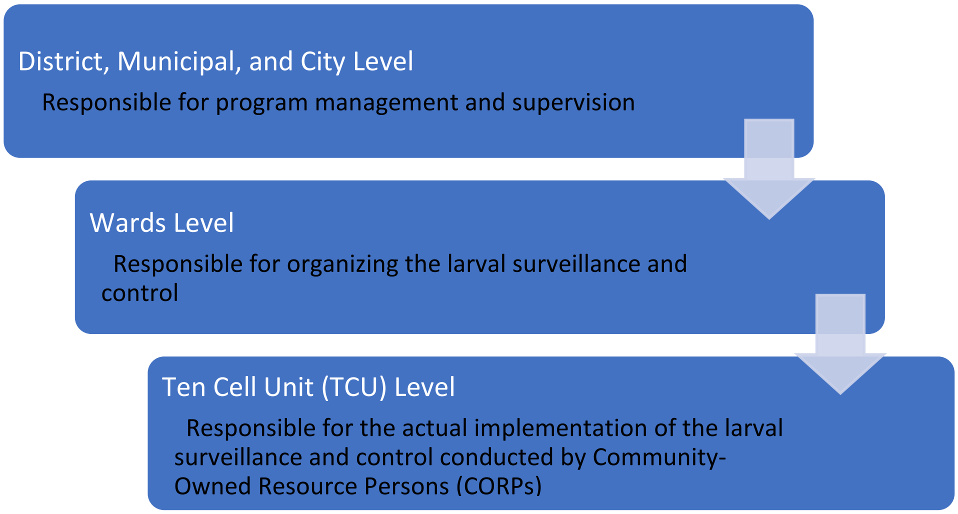

**Figure 1.** The Urban Malaria Control Program (UMCP) in Tanzania produced many studies on community-based vector surveillance and control in Africa in this scoping review. A case study of the UMCP program is presented here and the management structure which facilitated the community-based urban larval surveillance and control activities is described emphasizing the municipality, district, and city leadership.

All the programs implemented by the UMCP were conducted by community-owned resource persons (CORPs), whom were community members chosen by street health committees. CORPs were responsible for a specific section of ten-cell units (TCUs) and were compensated a minimal amount as part-time employees through the employment system developed for sundry small scale maintenance tasks (Vanek et al. 2006). The main challenges that the UMCP initially faced included participant concerns of working environment, workload, payment, and access issues for fenced or private plots of land. All of those challenges were better understood through a cross-sectional survey and addressed through redistribution of TCUs for equitable workloads and increased interaction with homeowners (Chaki et al. 2009, 2011, 2014). Many institutional lessons were learned including that pilot programs should start on a small and manageable scale and build on experience with the pilot programs having clearly designed plans that delineate who will do what, at what spatial scale, and how institutions should interact. Lastly, the capacity to manage the logistics, human resources, institutional partnerships, and funding support is just as important as the entomological skill present (Chaki et al. 2014). Many program development lessons were also learned from the UMCP with importance placed on community ownership, recruitment processes, government involvement & integration, geographic & social boundaries, and sustainability (Chaki et al. 2011, 2014, Fillinger et al. 2008). The main recommendations that should be gathered are to have community leaders conduct recruiting, to recruit participants from existing vector control programs, and to focus on incentives and participant motivation to increase community ownership (Chaki et al. 2011, Fillinger et al. 2008). The UMCP increased participant motivation by promoting the best performing CORPs to ward level supervisors and by offering full-time technician jobs to those CORPs who excelled at taxonomy (Chaki et al. 2014). During the beginning steps is also when consideration should be placed on the type of compensation that will be given to participants, to ensure it is sustainable for the government and that it will be motivating enough for participants to continue working. It is critical to involve the government in all steps from the start to ensure that all decisions made are realistic for the government and for early and effective integration into vertical systems (Chaki et al. 2014). The UMCP successfully involved the government by training current government health officers in basic entomology to conduct quality control checks of the CORPs and report summarized findings to higher level authorities. Lastly, it is critical to consider both the geographic and social boundaries when delegating work areas and workloads to increase ownership and avoid conflict (Chaki et al. 2014). The UMCP avoided this issue by recruiting participants from within the areas they would work which increased access to the local knowledge that participants bring with them in terms of potential breeding habitats and social contexts, reduced time spent travelling to work, and ensured acceptance from fellow neighbors (Chaki et al. 2011, Dongus et al. 2007). This community-based pilot work by the UMCP has been demonstrated and the model has now been translated at national scale through the National Malaria Control Program (NMCP). The NMCP in Tanzania now implements the community LSM method in 63 districts in the country. For malaria surveillance and control overall, a system that mobilizes and involves community members, is integrated into current vertical systems, and managed by either local or municipal health centers is recommended given its engagement of varying levels of stakeholders and potential for sustainability (Chaki et al. 2014). Additionally, results from the study conducted by Maheu-Giroux found larviciding to be a cost-effective form of vector control in most transmission settings in Africa where the incidence is above 110-116 infections per 1,000 per year. The literature suggests that larviciding should be considered as part of an IVM approach, if local conditions are optimal for its use, especially in urban areas (Maheu-Giroux and

Castro, 2014). The UMCP show that larviciding can be a valuable vector control method when utilized supplementary to current control methods and is a vital part of an IVM package (Geissbuhler et al. 2009).
