## supplemental figure 1 for "Community-Based Entomological Surveillance and Control of Vector-Borne Diseases: A Scoping Review"

**Supplemental Figure 1: Search Queries**

**Search Query:** Explore the concept of community-based entomological surveillance in the field of entomological monitoring for vector-borne diseases in evaluating the impact and guiding the selection of vector control programs. To look at community-based entomological surveillance compared to conventional entomological surveillance methods and the health metrics used in both.

**Key words:** community-based entomological surveillance, vector-borne disease, vector control, IRS, ITNs, LLINs, bed nets, community, surveillance, entomology, entomological monitoring, collection, decentralized, centralized, longitudinal, mosquito-borne, tick-borne, conventional, arbovirus, arboviral, arthropod, community-based control, citizen science, science, gold standard, comparison

**Medline (OVID)**

(community* ADJ5 surveillance) AND (entomological OR mosquito* OR tick* OR arthropod* OR vector-borne* OR vector control)
1990 - ; English

**Embase (OVID)**

(community* ADJ5 surveillance) AND (entomological OR mosquito* OR tick* OR arthropod* OR vector-borne* OR vector control)
1990 - ; English ; Not pubmed/medline

**Global Health (OVID)**

(community* ADJ5 surveillance) AND (entomological OR mosquito* OR tick* OR arthropod* OR vector-borne* OR vector control)
1990 - ; English

**Scopus**

TITLE-ABS-KEY((community* W/5 surveillance) AND (entomological OR mosquito* OR tick* OR arthropod* OR vector-borne* OR "vector control"))
