## supplemental figure 2 for "Community-Based Entomological Surveillance and Control of Vector-Borne Diseases: A Scoping Review"

**Supplemental Figure 2: Data Extraction Template**

| **General Information** |
| --- |
| Study ID |
| Title |
| Lead author contact details |
| Region:<br>*Asia*<br>*Africa*<br>*North America*<br>*South America*<br>*Central America*<br>*The Caribbean*<br>*Europe*<br>*Oceania*<br>*Worldwide* |
| Country |
| District |
| Study site coordinates / name |
| Disease |
| Vector |
| Notes:<br>*If excluding, enter in the reason* |
| **Characteristics of Included Studies** |
| <u>Methods</u> |
| Methods used:<br>*Aspiration*<br>*Human Landing Catch*<br>*Oviposition Trap*<br>*Larvicide Application*<br>*CDC Light Trap*<br>*Larval Surveys*<br>*Tent Traps*<br>*Tick Drags*<br>*Mobile phone based*<br>*Door-to-door infestation surveys (manual inspection for vectors or feces)*<br>*Sensor boxes*<br>*Household Bug Notification (HN)*<br>*Sticky Traps (ST)*<br>*Other* |
| Aim of study / study question |
| Health Metric Used |
| Study Design:<br>*Randomized Controlled Trial*<br>*Non-randomized Experimental Study*<br>*Cohort Study*<br>*Cross Sectional Study*<br>*Case Control Study*<br>*Systematic Review*<br>*Qualitative Research*<br>*Prevalence Study* |

| |
|---|
| *Case Series*<br>*Case Report*<br>*Diagnostic Test Accuracy Study*<br>*Clinical Prediction Rule*<br>*Economic Evaluation*<br>*Text and Opinion*<br>*Other* |
| Start Date |
| End Date |
| Study Details |
| Focus:<br>*Comparison*<br>*Citizen Science*<br>*Recommendations*<br>*Community* |
| If comparison: what conventional method is used? |
| If comparison: what community-based method is used? |
| If citizen science: what method is used? |
| If recommendations: what method was used and what recommendation or direction is given? |
| If community: how was the community involved? |
| Limitations to chosen focus |
| Advantages to chosen focus |
| Did they include information / conclusions on cost-effectiveness of community-based methods:<br>*Yes*<br>*No* |
| If yes, what was the conclusion? |
| Overall Conclusions |
| Overall Recommendations |
