## supplemental table 1 for "Community-Based Entomological Surveillance and Control of Vector-Borne Diseases: A Scoping Review"

**Supplemental Table 1: Studies included in final analysis categorized by region of the world and country (n=107)**

***some studies are conducted in more than one country**

| | Chagas Disease | Dengue | Malaria | TBD | MBD[1] | *Total* |
|---|---|---|---|---|---|---|
| ***South America*** | | | | | | |
| *Argentina* | 9 | 0 | 0 | 0 | 0 | *9* |
| *Brazil* | 4 | 0 | 0 | 0 | 0 | *4* |
| *Bolivia* | 2 | 0 | 0 | 0 | 0 | *2* |
| *Colombia* | 3 | 0 | 0 | 0 | 0 | *3* |
| *Paraguay* | 0 | 1 | 0 | 0 | 0 | *1* |
| ***North America*** | | | | | | |
| *United States of America* | 1 | 0 | 0 | 4 | 2 | *7* |
| *Canada* | 0 | 0 | 0 | 3 | 0 | *3* |
| *Cuba* | 0 | 3 | 0 | 0 | 0 | *3* |
| *Mexico* | 1 | 0 | 0 | 0 | 0 | *1* |
| ***Central America*** | | | | | | |
| *Guatemala* | 3 | 0 | 0 | 0 | 1 | *4* |
| *Honduras* | 2 | 0 | 0 | 0 | 1 | *3* |
| *El Salvador* | 1 | 0 | 0 | 0 | 0 | *1* |
| *Nicaragua* | 1 | 0 | 0 | 0 | 0 | *1* |
| ***Africa*** | | | | | | |
| *Tanzania* | 0 | 0 | 7 | 0 | 1 | *8* |
| *Rwanda* | 0 | 0 | 5 | 0 | 0 | *5* |
| *Ethiopia* | 0 | 0 | 3 | 0 | 0 | *3* |
| *Kenya* | 0 | 0 | 2 | 0 | 0 | *2* |
| *Senegal* | 0 | 0 | 0 | 0 | 2 | *2* |
| *Malawi* | 0 | 0 | 1 | 0 | 0 | *1* |
| *Uganda* | 0 | 0 | 1 | 0 | 0 | *1* |
| *West Africa* | 0 | 0 | 0 | 0 | 1 | *1* |
| *Zambia* | 0 | 0 | 1 | 0 | 0 | *1* |
| *Burkina Faso* | 0 | 0 | 0 | 0 | 1 | *1* |
| *Benin* | 0 | 0 | 0 | 0 | 1 | *1* |
| *Cameroon* | 0 | 0 | 0 | 0 | 1 | *1* |
| *Cape Verde* | 0 | 0 | 0 | 0 | 1 | *1* |
| *Chad* | 0 | 0 | 0 | 0 | 1 | *1* |
| *Cote D'Ivoire* | 0 | 0 | 0 | 0 | 1 | *1* |

| | | | | | | |
|---|---|---|---|---|---|---|
| *Equatorial Guinea* | 0 | 0 | 0 | 0 | 1 | *1* |
| *Gabon* | 0 | 0 | 0 | 0 | 1 | *1* |
| *Gambia* | 0 | 0 | 0 | 0 | 1 | *1* |
| *Ghana* | 0 | 0 | 0 | 0 | 1 | *1* |
| *Guinea* | 0 | 0 | 0 | 0 | 1 | *1* |
| *Guinea Bissau* | 0 | 0 | 0 | 0 | 1 | *1* |
| *Liberia* | 0 | 0 | 0 | 0 | 1 | *1* |
| *Mali* | 0 | 0 | 0 | 0 | 1 | *1* |
| *Mauritania* | 0 | 0 | 0 | 0 | 1 | *1* |
| *Niger* | 0 | 0 | 0 | 0 | 1 | *1* |
| *Nigeria* | 0 | 0 | 0 | 0 | 1 | *1* |
| *Sao Tome and Principe* | 0 | 0 | 0 | 0 | 1 | *1* |
| *Sierra Leone* | 0 | 0 | 0 | 0 | 1 | *1* |
| *Togo* | 0 | 0 | 0 | 0 | 1 | *1* |
| ***Asia*** | | | | | | |
| *India* | 0 | 1 | 0 | 0 | 1 | *2* |
| *Indonesia* | 0 | 1 | 0 | 0 | 0 | *1* |
| *Myanmar* | 0 | 1 | 0 | 0 | 0 | *1* |
| *Thailand* | 0 | 1 | 0 | 0 | 0 | *1* |
| *Vietnam* | 0 | 1 | 0 | 0 | 0 | *1* |
| ***Europe*** | | | | | | |
| *Germany* | 0 | 0 | 0 | 0 | 2 | *2* |
| *The Netherlands* | 0 | 0 | 0 | 1 | 1 | *2* |
| *Spain* | 0 | 0 | 0 | 0 | 2 | *2* |
| *France* | 0 | 0 | 0 | 0 | 1 | *1* |
| *Hungary* | 0 | 0 | 0 | 1 | 0 | *1* |
| *Portugal* | 0 | 0 | 0 | 0 | 1 | *1* |
| *United Kingdom* | 0 | 0 | 0 | 0 | 1 | *1* |
| ***Oceania*** | | | | | | |
| *Australia* | 0 | 0 | 0 | 0 | 2 | *2* |
| *Solomon Islands* | 0 | 0 | 0 | 0 | 1 | *1* |
| *Vanuatu* | 0 | 1 | 0 | 0 | 0 | *1* |
| ***Caribbean*** | | | | | | |
| *Haiti* | 0 | 0 | 1 | 0 | 0 | *1* |
| ***Total*** | **27** | **10** | **21** | **9** | **40** | ***107*** |

[1]MBD = all other articles focusing on MBDs exclusive of dengue or malaria
