## supplemental table 2 for "Community-Based Entomological Surveillance and Control of Vector-Borne Diseases: A Scoping Review"

**Supplemental Table 2: Breakdown of Capture Method – "Other" by Disease**

| | Chagas Disease | Dengue | Malaria | TBD | MBD[1] | General VBD | ***Total*** |
|---|---|---|---|---|---|---|---|
| *[2]Home Improvements* | 0 | 0 | 1 | 0 | 0 | 0 | **1** |
| *[2]Mesocyclops Application* | 0 | 1 | 0 | 0 | 0 | 0 | **1** |
| *Passive Tick Collection* | 0 | 0 | 0 | 8 | 0 | 0 | **8** |
| *Passive Mosquito Collection* | 0 | 0 | 1 | 0 | 5 | 0 | **6** |
| *[2]Fumigant Canisters* | 1 | 0 | 0 | 0 | 0 | 0 | **1** |
| *BG Sentinel Trap* | 0 | 0 | 0 | 0 | 2 | 0 | **2** |
| *Handmade Carbon-Dioxide Baited Traps* | 0 | 0 | 1 | 0 | 0 | 0 | **1** |
| *M-Trap* | 0 | 0 | 0 | 0 | 1 | 0 | **1** |
| *Angulo Traps* | 1 | 0 | 0 | 0 | 0 | 0 | **1** |
| *BG Gravid Trap* | 0 | 0 | 0 | 0 | 1 | 0 | **1** |
| ***Total*** | **2** | **1** | **3** | **8** | **9** | **0** | **23** |

[1]MBD = all other articles focusing on MBDs exclusive of dengue or malaria

[2]Common vector control methods / methods commonly used across disease systems
